# Human cortical dynamics reflect graded contributions of local geometry and network topography

**DOI:** 10.1101/2025.01.07.631564

**Authors:** Jessica Royer, Casey Paquola, Raul Rodriguez-Cruces, Hans Auer, Alexander Ngo, Ella Sahlas, Thaera Arafat, Chifaou Abdallah, Daniel Mansilla de Latorre, Raluca Pana, Jeffrey Hall, Robert Leech, Jonathan Smallwood, Birgit Frauscher, Boris C. Bernhardt

**Author notes:** Corresponding authors: Jessica Royer, Psy.D., Ph.D. Boris C. Bernhardt, Ph.D.

## Abstract

The brain is a physically embedded and heavily interconnected system that expresses neural rhythms across multiple time scales. While these dynamics result from the complex interplay of local and inter-regional factors, the relative contribution of such mechanisms across the cortex remains unclear. Our study explores geometric, microstructural, and connectome-level constraints on cortex-wide neural activity. We leverage intracranial electroencephalography recordings to derive a coordinate system of human cortical dynamics. Using multimodal neuroimaging, we could then demonstrate that these patterns are largely explainable by geometric properties indexed by inter-regional distance. However, dynamics in transmodal association regions are additionally explainable by incorporation of inter-regional microstructural similarity and connectivity information. Our findings are generally consistent when cross-referencing electroencephalography and imaging data from large-scale atlases and when using data obtained in the same individuals, suggesting subject-specificity and population-level generalizability. Together, our results suggest that the relative contribution of local and macroscale constraints on cortical dynamics varies systematically across the cortical sheet, specifically highlighting the role of transmodal networks in inter-regional cortical coordination.

## Introduction

The brain generates nested and propagating rhythms from the synchronized firing of neuronal ensembles (*1*). These dynamic signals are a core feature of brain organization, serving as established markers of brain health (*2–4*), neural communication (*5–7*), as well as vigilance and cognitive states (*8–15*). Intracranial electroencephalography (iEEG) offers a unique window into these rhythms. Moving beyond spatially restricted recordings performed in individual patients (*16, 17*), recent data sharing initiatives have provided comprehensive maps of human neurophysiology and unlocked new avenues to study inter-regional heterogeneity in neural dynamics (*8, 18*). However, technical challenges relating to multimodal data availability, registration, and integration across and within patient cohorts have particularly hindered these investigations, which have so far been limited to single scales of inquiry. The present work fills this gap by uncovering large-scale gradients of neural dynamics and relating this complex activity landscape to underlying multiscale anatomy.

Prior work has demonstrated distinct neurophysiological signatures operating across distributed cortical territories (*19*). These signatures generally follow overlapping spatial gradients of cortical microarchitecture involving neurotransmitter receptor and transporters, gene expression, metabolic demands, cortical thickness, and proxies of intracortical myelination (*20–22*). Relatedly, focal lesions altering local anatomy and microarchitecture co-localize with atypical neural dynamics (*2, 3*). While this evidence supports the role of intrinsic physiological mechanisms in generating regional neural activity (*23, 24*), local structural features can only partially explain spatial variations in neural dynamics. For instance, areas made up of fundamentally different milieus can also produce rhythms with similar features: peaks in beta-range activity have been recorded in large parts of the frontal lobe, from the precentral gyrus to the frontal pole, as well as portions of the parietal lobe and the anterior insula (*18*). As such, network-level propagation of brain activity via direct structural connections as well as polysynaptic functional interactions has also been suggested to account for this inter-regional variability (*25*). Indeed, local dynamics systematically vary along the cortical processing hierarchy, as demonstrated by task paradigms, stimulation work, and *in silico* explorations (*26–28*). Resting dynamics also propagate to nearby regions via travelling waves, scaffolded by the brain’s short- and long-range connections (*29*), producing a complex mosaic of intrinsic and extrinsic constraints on neural dynamics.

Such multiscale inquiries can be achieved by integrating the high spatiotemporal precision of iEEG with recent methodological and analytical advances in magnetic resonance imaging (MRI) offering complementary insights into cortico-cortical network organization. At the most basic level is cortical geometry, which reflects the proximity of brain regions. The brain is a physically embedded network, with adjacent regions generally participating in similar functional processes (*30*). Moreover, adjacent regions are generally strongly connected via intracortical axon collaterals and superficial white matter fibres (*31*), which closely regulate local neural activity (*32, 33*). A further important component of brain network architecture is local intracortical microstructure, which has classically been studied using cytoarchitectonic mapping of *post mortem* tissue (*34*). Recently, similar approaches have been translated to *in vivo* MRI using myelin-sensitive contrasts, such as T1 relaxometry (*35–38*). Both lines of investigation suggest that regions with similar microstructure generally participate in corresponding functional processes and are more likely to share a direct structural connection (*34, 35, 39, 40*). Beyond the study of local geometry and microstructure, techniques such as diffusion MRI tractography and functional MRI connectivity have characterized large-scale, regionally distributed structural and functional networks. While the former approximates long-range, white matter fibre tracts *in vivo* (*41–44*), the latter taps into spatially distributed, and often polysynaptic intrinsic networks (*45–47*). In line with recently developed composite wiring diagrams of the human cortex emphasizing a synergy of different MRI modalities in predicting macroscale function (*48–54*), simultaneous consideration of these diverse modalities elucidates the contribution of short- and long-range network information to human cortical dynamics.

Our goal was *(i)* to chart the cortical topography of neural rhythms in the human brain interrogated with iEEG (*18*), and *(ii)* identify the differential association between cortical dynamics and network organization mapped with multimodal MRI (*55*). Applying dimensionality reduction to a cortex-wide assessment of inter-regional iEEG power spectra similarity allowed us to situate cortical areas within a compact embedding space constrained by their dynamics. These gradients, mapped here in iEEG data, captured the coordinate system of neural rhythms in the neocortex, grouping regions with similar neurophysiology independently of their anatomical location. We then quantified the contribution of different network level features to the topography of neural dynamics using multilinear spatial statistics at a cortex-wide scale. We complemented this approach with region-specific models capturing the relative contribution of each network metric across cortical areas, and tracked how these contributions varied along the cortical processing hierarchy (*56*). Our assessment was based on independent datasets and systematic variations of analysis parameters to identify robust associations between human cortical dynamics and multiscale network features.

## Results

### Mapping cortical gradients of neural dynamics

For the main analyses, we leveraged a Discovery cohort (MNI open iEEG atlas, n=106) in which iEEG data were aggregated in stereotactic space (*18*). We assessed correspondence to neural organization derived from two high-definition MRI datasets acquired in 50 healthy individuals at 3T and 10 healthy individuals at 7T. Our main analyses thus rest on a design comparing iEEG and imaging modalities across two different datasets. We then evaluate the generalizability of our findings using a fully independent Replication cohort (MNI clinical dataset, n=18), where both iEEG and multimodal pre-surgical MRI data were available in each participant for within-cohort comparison of modalities.

The MNI open iEEG atlas includes recordings from 106 patients with intractable epilepsy, totalling over 1,700 data points taken from non-epileptic channels registered to the MNI152 reference space (*18*). Mapping all channels to a single hemisphere as in previous work (*8, 18*) granted wide sampling of the cortical sheet (**Figure 1A**). Spatial heterogeneity of spectral features was visible across channels, but could also be recovered when averaging channel power spectra within larger cortical regions (**Figure 1B**) (*57*). We captured spatial variations in macroscale neural dynamics by applying dimensionality reduction to similarity patterns in regional power spectra (**Figure 1C**). After averaging vertex-wise, log-transformed power spectral densities (PSDs) within parcels, regional PSDs were cross-correlated while accounting for the cortex-wide average power spectrum using partial correlations. We converted the result to a normalized angle affinity matrix, and applied diffusion map embedding, a non-linear dimensionality reduction technique, to recover eigenvectors (*gradients*) of macroscale similarity in regional neural dynamics. We retained gradients explaining a cumulative sum of 50% of the variance for further analyses (G1-G14; see **Figure S1A** for the topography of these gradients). The embedding space formed by the first two gradients (explaining 17.37% of the variance in PSD similarity) was used as a complementary visualization of findings.

**Figure 1.**
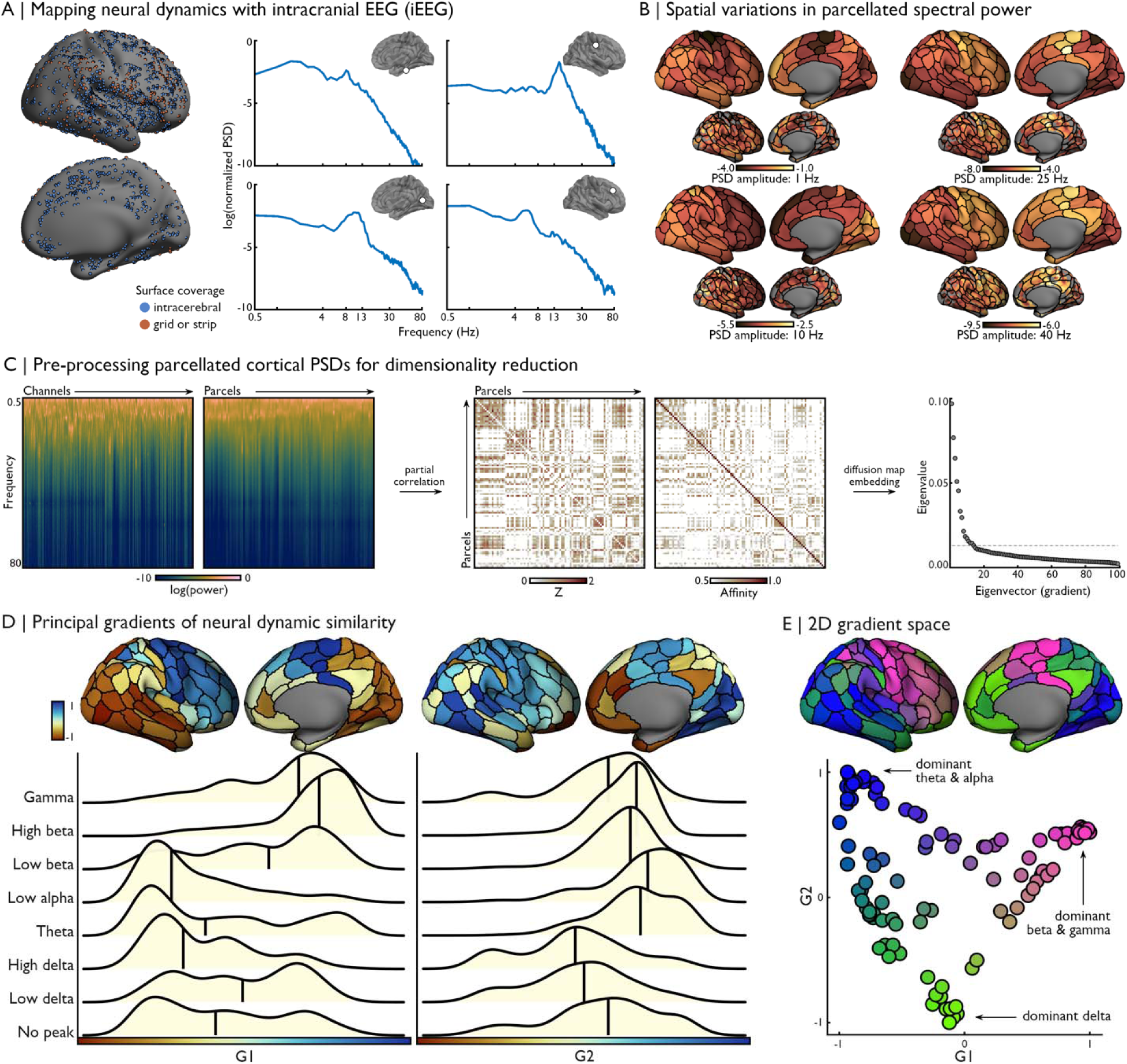
Cortical gradients of neural dynamics. **(A)** Channel coordinates were registered to the fsLR-32k surface template. The power spectral density (PSD) of each channel was computed and normalized as in previous work, and is plotted in log-log space (*18*). **(B)** Channel PSDs were mapped to single vertices, expanded to 5mm regions surrounding each assigned vertex (channel region), and averaged within parcels. PSD amplitude at selected frequencies (1-10-25-40Hz) mapped to channel regions (bottom) and the Schaefer-200 parcellation (top) highlight the spatial heterogeneity of neural dynamics across the cortex. **(C)** After averaging channel-wise PSDs within parcels, features were cross-correlated to generate macroscale gradients of neural rhythm similarity using diffusion map embedding. The dashed line marks the 14^th^ gradient, at which the cumulative sum of explained variance by each gradient reached 50%. **(D)** Projection of G1 and G2 to the cortical surface emphasized strong differentiation of neural dynamics between motor-sensory (G1; blue-red) and unimodal-transmodal (G2; blue-red) regions. Ridge plots show how each gradient captured different spatial axes of variance in neural dynamics, leveraging previously computed frequency-band classes for this dataset (*18*). **(E)** The scatter plot depicts node positions in the 2D gradient space, coloured according to proximity to axis limits, alongside an equivalent representation on the cortical surface. Source data are provided as a Source Data file.

The first gradient (G1) evolved along the anterior-posterior direction, mainly differentiating primary motor and surrounding lateral prefrontal cortices from unimodal sensory regions (**Figure 1D**). G1 indeed mainly followed the structural stratification defined by Von Economo and Koskinas (*58–60*). This suggests a close correspondence between the axis of neurophysiological differentiation captured by G1 and regional heterogeneity in laminar differentiation, notably segregating “input” sensory regions with a granular structure from “output” motor areas with an agranular structure (**Figure S1B**). This gradient mirrored regional variations in spectral peaks, by segregating areas with strong beta and gamma-range activity from areas showing prominent delta, theta, and alpha-range peaks (*18*). Next, G2 differentiated unimodal sensorimotor from transmodal cortices, recapitulating the previously established principal axis of cortical connectivity and functional hierarchy (**Figure 1D**) (*61*). Indeed, G2 was specifically correlated with the unimodal-transmodal gradient of cortical organization derived from functional MRI (Spearman *r*=0.564; *p*_spin_<0.001; **FIGURE S1C; correlations with all other gradients did not survive Bonferroni correction**). This pattern segregated regions showing predominant lower frequency activity, from other cortical areas with peaks occurring above the delta frequency range. The two-dimensional representation of neural dynamics formed by these gradients separated unimodal sensory regions, precentral motor areas, and spatially distributed transmodal systems (**FIGURE 1E**). Cortical surface plot shown in **FIGURE 1B** furthermore highlighted consistency of regional power spectra amplitudes with previously published normative maps of neocortical dynamics (*2, 3, 62*). We observed higher delta band power in frontal, parietal, and temporal cortices relative to unimodal sensory and motor regions. High alpha- and beta-range power was observed within occipital and precentral regions, respectively, while a rostro-caudal shift was observed in gamma-range activity across the cortex. Collectively, these result map gradients of neural rhythms able to resolve fine-grained oscillatory activity of individual channels. We additionally show that these gradient recover macroscale trends in the overall similarity of spectral content across the cortex.

A series of control analyses were implemented to assess the robustness of the described gradients and embedding space. First, results were robust to the use of different parcellations schemes (**Figure S2**). Indeed, generating gradients from unparcellated PSD data (*i.e.*, channel regions – see **Materials and Methods**) or averaging PSDs within nodes defined by two different atlases (*i.e.,* Glasser-360 and a subdivision of the aparc parcellation) yielded similar topographies for both G1 (vertexwise Spearman correlations for unparcellated: r=0.673; Glasser-360: r=0.839; aparc-200 r=0.864) and G2 (vertexwise Spearman correlations for unparcellated: r=0.524 ; Glasser-360: r=0.580; aparc-200 r=0.645). Furthermore, we could recover the topography of the first 10 gradients when imposing an upper bound to the number of channels included in each parcel of the Schaefer-200 parcellation (**Figure S3**), suggesting that our findings could not be attributed to variable channel coverage across nodes. We also replicated our findings when preserving the bilateral distribution of channels and applying a lower resolution parcellation (Schaefer-100) to maintain broad cortical coverage (**Figure S4**). The high correspondence of G1 (vertexwise Spearman correlation r_left_=0.767; r_right_=0.804) and G2 (vertexwise Spearman correlation r_left_=0.645; r_right_=0.671) topographies with gradients derived from our main analysis provides additional support for the independence of our findings to parcellation choice. Lastly, spatial correlations assessing the consistency of gradients across a range of hyperparameter combinations were generally very strong (**Figure S7**).

### Multiscale connectomics of neural dynamics

After establishing the topography of macroscale gradients of neural dynamics, we investigated how this embedding space co-varied with multiscale properties of brain network organization. All network measures were gathered from an independent cohort of 50 unrelated, healthy volunteers that underwent multimodal 3T MRI (23 women; age mean ± standard deviation = 30.28 ± 5.74 years) (*55*). We first computed the Euclidean distance between all node pairs in the embedding space, which served as a proxy for inter-node similarity in neural dynamics (*embedding distance*, **Figure 2A**). This matrix was highly robust to the number of gradients included when calculating inter-nodes distances (**Figure S8**). Node proximity in the embedding space was cross-referenced with measures of *(i)* cortical geometry indexed by inter-node geodesic distance computed along the surface mesh (geodesic distance, GD), *(ii)* tractography-based structural connectivity (SC), *(iii)* intrinsic functional connectivity generated from resting-state functional MRI (rs-fMRI), and *(iv)* microstructural profile covariance (MPC) derived from T1 relaxometry. Matrices were computed for each participant and averaged across individuals (for details on data processing, analysis, and group-level aggregation, see **Materials and Methods**). Multimodal associations were computed by correlating the upper triangular edges of each matrix. We only considered MRI data from the right hemisphere, as all iEEG data had been transposed to the right hemisphere to improve cortical coverage. For each modality, statistical significance was determined by comparing empirically observed correlations to those obtained with surrogate matrices in which each row had been shuffled using spatial autocorrelation-preserving null models (*63, 64*).

**Figure 2.**
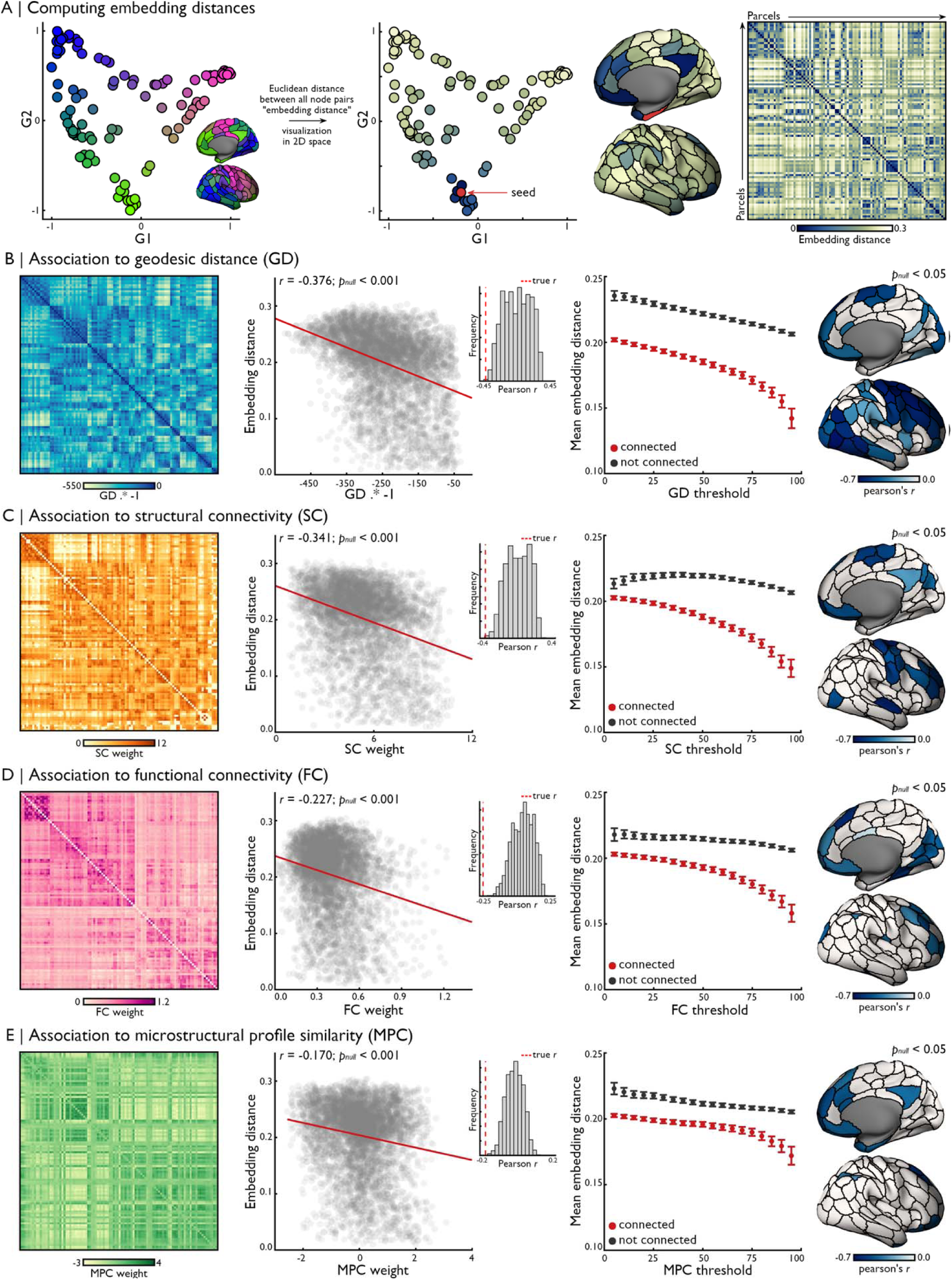
Multiscale network correlates of cortical neural dynamics. **(A)** The Euclidean distance between all node pairs (*embedding distance*) in the gradient space (here, composed of G1-G14) reflected the similarity of neural dynamics between every node pair, with an example shown for a seed in the mesiotemporal lobe. Data is visualized using a 2-dimensional embedding space (G1/G2) and along the cortical surface. Significant correlations were found between upper triangle edges of the embedding distance matrix and **(B)** geodesic distance (multiplied by -1 to facilitate interpretation and comparison with other network metrics; n=4,950 edges), **(C)** tractography-based SC (n=4,548 edges), **(D)** FC computed from rs-fMRI (n=4,947 edges), **(E)** and MPC derived from T1 relaxometry (n=4,950 edges). Number of analysed edges differ for SC and FC because of the exclusion of edges with a value of 0 (for both SC and FC) and negative values (only FC). Red line indicates least-squares regression fit. Statistical significance was determined by comparing empirically observed correlations to those obtained with surrogate matrices in which each row had been shuffled using spatial autocorrelation-preserving null models (1,000 permutations) (*63, 64*). The distribution of correlations obtained from these surrogate matrices are displayed in the inset histogram for panels B-E, in which the dashed red line represents the true correlation obtained in the data. Analogous analyses using binarized network matrices (right) yielded consistent results across a wide range of thresholds. For each threshold *i*, edges were labelled as “connected” if their weight was within the *i%* highest values for the analysed modality (total of 19 thresholds tested on the number of edges included in the associated scatter plot for each modality). Corresponding embedding distances at each edge labelled “connected” were then averaged, and compared to the average embedding distances of edges labelled “not connected”. Error bars denote the standard error of the mean. Node-level correlations are displayed on the cortical surface. Source data are provided as a Source Data file.

We first observed a moderate correlation (*r*=-0.376; *p_null_*<0.001) between GD of regions along the cortical sheet and embedding distance, indicating that nodes with similar dynamics were situated closer to each other along the cortical sheet (**Figure 2B**). Node-wise correlations between both modalities were particularly strong in anterior and posterior lateral regions. We also obtained a moderate correlation (*r*=-0.341; *p_null_*<0.001) between edge strength of the SC matrix and embedding distance, suggesting closer neural dynamic similarity between regions sharing stronger structural connections (**Figure 2C**). Regional SC and embedding distances were most strongly correlated in lateral frontal and temporal regions. Polysynaptic network architectures captured by the functional connectome were weakly, yet significantly, correlated with embedding distances (*r*=-0.227; *p_null_*<0.001), once again indicating stronger connections between regions sharing similar neural dynamics (**Figure 2D**). This relationship was particularly strong in medial and lateral prefrontal, as well as medial occipito-temporal regions. Lastly, embedding distance was significantly yet more weakly correlated with MPC (*r*=-0.170; *p_null_*<0.001), with significant correlations confined to heteromodal and paralimbic cortical structures such as mesial temporal and frontal regions, as well as the precuneus and lateral parietal areas (**Figure 2E**). Of note, relationships between neocortical dynamic similarity and multiscale connectivity measures were consistent when binarizing network matrices across a wide range of thresholds (**Figure 2B-E**). Findings were replicable when cross-referencing embedding distances with multimodal data collected in a distinct set of participants scanned at 7T MRI (*65*) (**Figure S5**), and when computing correlations between embedding distances and MRI measures from bilaterally mapped data (SC *r*=-0.322; FC *r*=-0.217; MPC *r*=-0.185). Together, these results demonstrate that neural dynamics similarity is most strongly associated with cortical geometry indexed by geodesic distance, with weaker yet statistically significant relationships to large-scale network metrics. Modality-specific correlations were nonetheless highly variable across regions.

These results show regional variations in the relative contributions of different network properties to cortical dynamics, while emphasizing the stronger relationship with spatial proximity between nodes. We next sought to clarify the combined contribution of network features to neural dynamic similarity by quantifying potential gains in model accuracy when considering multiscale network properties over GD alone. To do so, we constructed several multiple linear regression models, in which we varied the combination of predictors implemented in the model as well as the nature of the predicted and predictor variables. A baseline model including GD as its only predictor was used to benchmark the performance of the more complex models incorporating multimodal predictor sets. These models encompassed partial models with two (GD and another modality) or three predictors (GD and two other modalities) in addition to a full model with four predictors (all modalities; **Figure 3A**).

**Figure 3.**
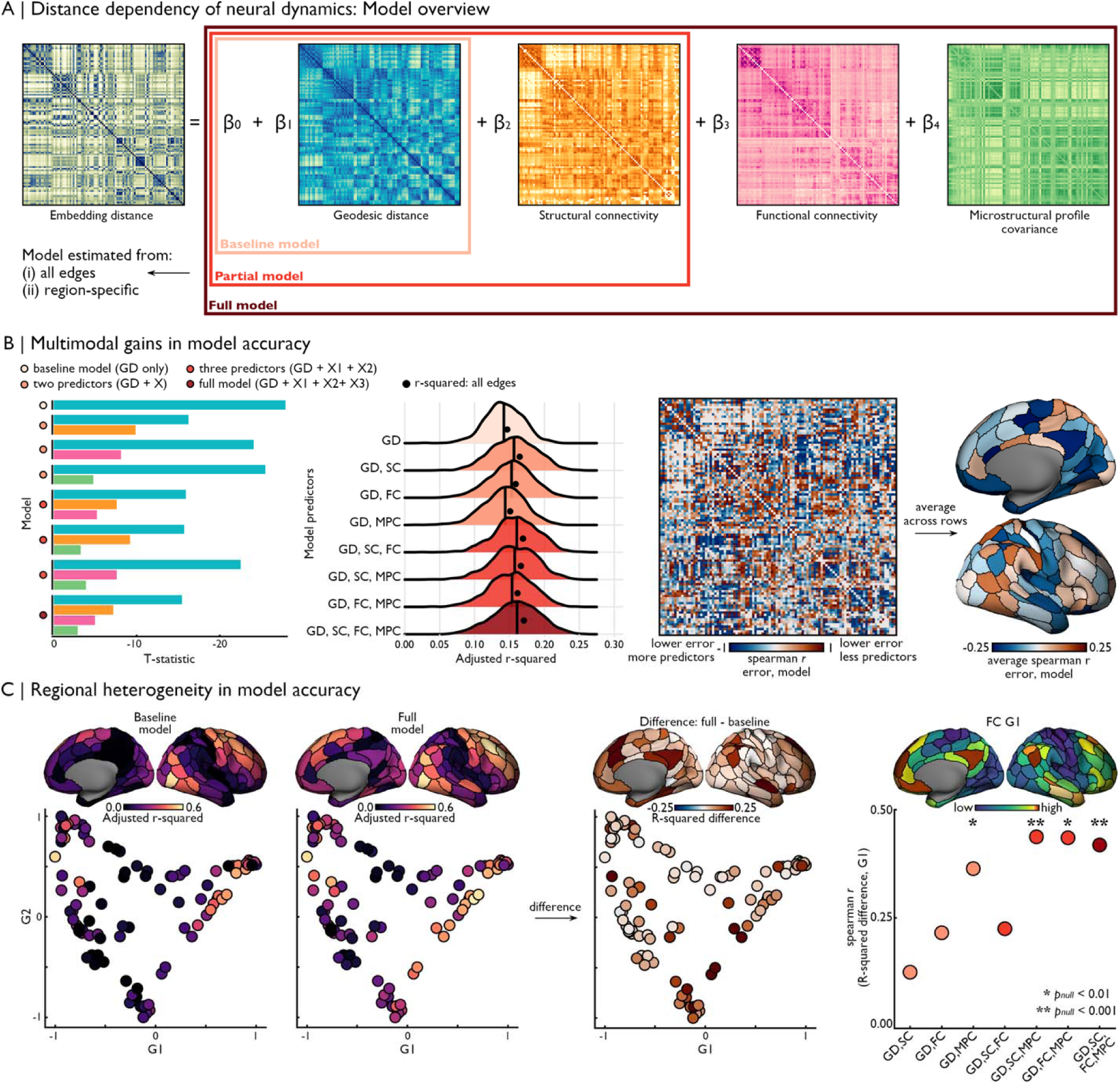
Unique and shared contributions of multiscale network properties to neural dynamics. **(A)** A series of multiple linear regression models integrating different sets of predictors were constructed to assess the association between multiscale network parameters and neural dynamics. A baseline model including GD as its only predictor was used to benchmark the performance of the more complex models leveraging multimodal predictor sets. **(B)** Bar plots of T-statistics for each predictor in each model showed that GD was the strongest contributor to model accuracy when constructing models across all edges. The colour of the bars corresponds to the different MRI modalities shown in panel A (blue: GD; orange: SC; pink: FC; green: MPC). Coloured dots along the *y*-axis denote model category as shown in panel A. Ridge plots show that variance explained (adjusted r^2^) was stable across models. Each ridge plot shows the distribution of bootstrap results, with their median value indicated by the black line. The black dot marker indicates the true adjusted r^2^ for each model. We assessed spatial patterns of region-specific gains in model accuracy by model complexity, indexed by correlating the number of model predictors with the mean squared error produced at each edge for the corresponding model. Averaging resulting spearman *r* values across edges for each node (*i.e.,* across rows) showed that adding predictors resulted in variable accuracy gains across the cortex (*blue*: higher accuracy with more predictors; *red*: lower accuracy with more predictors). **(C)** Independently fitting multiple linear regression models for each node’s connections and comparing accuracies across models showed larger absolute gains in model accuracy in transmodal cortices (located at the apex of the cortical processing hierarchy represented by the principal gradient of intrinsic functional connectivity). Regional increases in model accuracy relative to the baseline model were only significantly correlated with the principal gradient for models including both GD and MPC. Statistically significant correlations are indicated by asterisks (\**p_nul_*_l_<0.01; \*\**p_nul_*_l_<0.001). *Abbreviations*: GD: Geodesic distance; SC: Structural connectivity; FC: Functional connectivity; MPC: Microstructural profile covariance; ED: Embedding distance. Source data are provided as a Source Data file.

Constructing this model across all unique edges (*i.e.*, the upper triangle of each matrix) highlighted the strong contribution of GD relative to other modalities across all models (**FIGURE 3B**). Notably, more complex models provided minimal gains in prediction accuracy, either when implementing the model on all data points (range of adjusted r^2^=0.147-0.171) or when bootstrapping the data (90% train and 10% test; 1000 permutations; range of median adjusted r^2^=0.142-0.161). We assessed cross-model accuracy variations by correlating the prediction error for each edge with the number of predictors in the corresponding model. In this approach, negative correlations indicate decreasing error with more complex models (*i.e.,* high prediction accuracy for models with more predictors), while positive correlations indicate increasing error with more complex models (*i.e.,* including additional predictors resulted in a noisier model). Our results show overall accuracy gains with increasing model complexity in mesial and anterolateral temporal, orbitofrontal, and dorsolateral prefrontal cortices, as well as the precuneus, while opposite trends were observed primarily in occipito-parietal regions (**FIGURE 3B, right**). These findings indicate heterogeneous contributions of network predictors across cortical regions.

We further explored region-specific associations between multiscale network features to neural dynamic similarity by fitting distinct multiple linear regression models for each node’s connections and comparing model accuracy across different predictor combinations (**Figure 3C**). For instance, when comparing baseline (GD only) and full (all network features) models, the greatest gains in adjusted r^2^ were seen in dorsal lateral and medial frontal cortices, orbitofrontal, mesiotemporal, lateral parietal and precuneus regions. Regional differences in accuracy gains across certain model pairs were significantly correlated with the cortical processing hierarchy represented by the principal gradient of intrinsic functional connectivity (*56, 66*). More specifically, regional gains in model accuracy were significantly correlated with the principal gradient in models including both GD and MPC as predictors over GD alone, and accuracy could be further increased by including either SC or FC. These findings were replicated using an alternate representation of this hierarchy mapping four functional zones co-localizing with regional changes in cortical laminar elaboration (*61, 67*) (**Figure S6**). These findings suggest a complementarity of local geometry and microstructure in supporting macroscale similarity of neural dynamics.

### Replicability of gradients of neural dynamics

Our work mapped the landscape of neural dynamics similarity of the cortex and decoded its multiscale network correlates using multilinear spatial statistics. We lastly leveraged an independent patient cohort to assess the replicability of macroscale gradients of neural dynamics and their association with within-sample neuroimaging metrics. The *MNI clinical dataset* (*Replication cohort*) includes 18 patients who underwent high-resolution multimodal MRI prior to clinical iEEG investigations, allowing for precise, patient-specific segmentation of cortical grey matter, co-registration of iEEG contacts, and cross-referencing to within-sample connectivity measurements. Although all data modalities were available in patients of the Replication cohort, these replication analyses study a completely distinct sample of participants relative to the Discovery cohort, for both iEEG and MRI metrics.

Following the same analytical pipeline as in our main analysis, electrophysiological data from all patients was combined to generate group-level gradients of neural dynamic similarity (**Figure 4A**). The gradient space in this independent sample segregated temporo-parietal regions and frontal cortices along G1, and distinguished transmodal temporal and frontal structures from lower-order sensorimotor areas along G2 (**Figure 4B**). The first 13 gradients, explaining a cumulative 50% of the variance in the input data, were all significantly correlated (*p_spin_*<0.05; *p*<0.003 with Bonferroni correction) with their corresponding gradient in the Discovery sample (after Procrustes alignment including translation, rotation, and scaling: mean±SD Spearman *r*=0.509±0.076). After alignment, we found a significant correlation between the embedding distance matrices of Discovery and Replication cohorts (*r*=0.304; *p*<0.001), with variable similarity across regions (**Figure S9**). We also applied a more conservative alignment strategy where Replication cohort gradients were reordered to optimally match (*i.e.,* minimize the distance) with the Discovery cohort eigenvectors. With this approach, correlations were overall lower but also indicated generalizability of main findings in the Replication cohort (mean±SD Spearman *r*=0.196±0.154; **Figure S10**). Systematic within-cohort and cross-cohort correlations spanning the gradient hyperparameter space found the resulting gradients to be generally robust to parameter selection (**Figure S7**). Differences in vertex-wise channel coverage between both cohorts partly contributed to node-wise displacement in the embedding space (**Figure 4B**). Channel coverage in the Replication cohort was notably higher in lateral temporal regions, and lower in insular and frontal cortices. The coverage difference map across datasets was significantly correlated with average node rank changes between each gradient (*r*=-0.179; *p_spin_*=0.038), suggesting that regional differences in channel coverage across the datasets may partially account for node shifts in the resulting embedding space.

**Figure 4.**
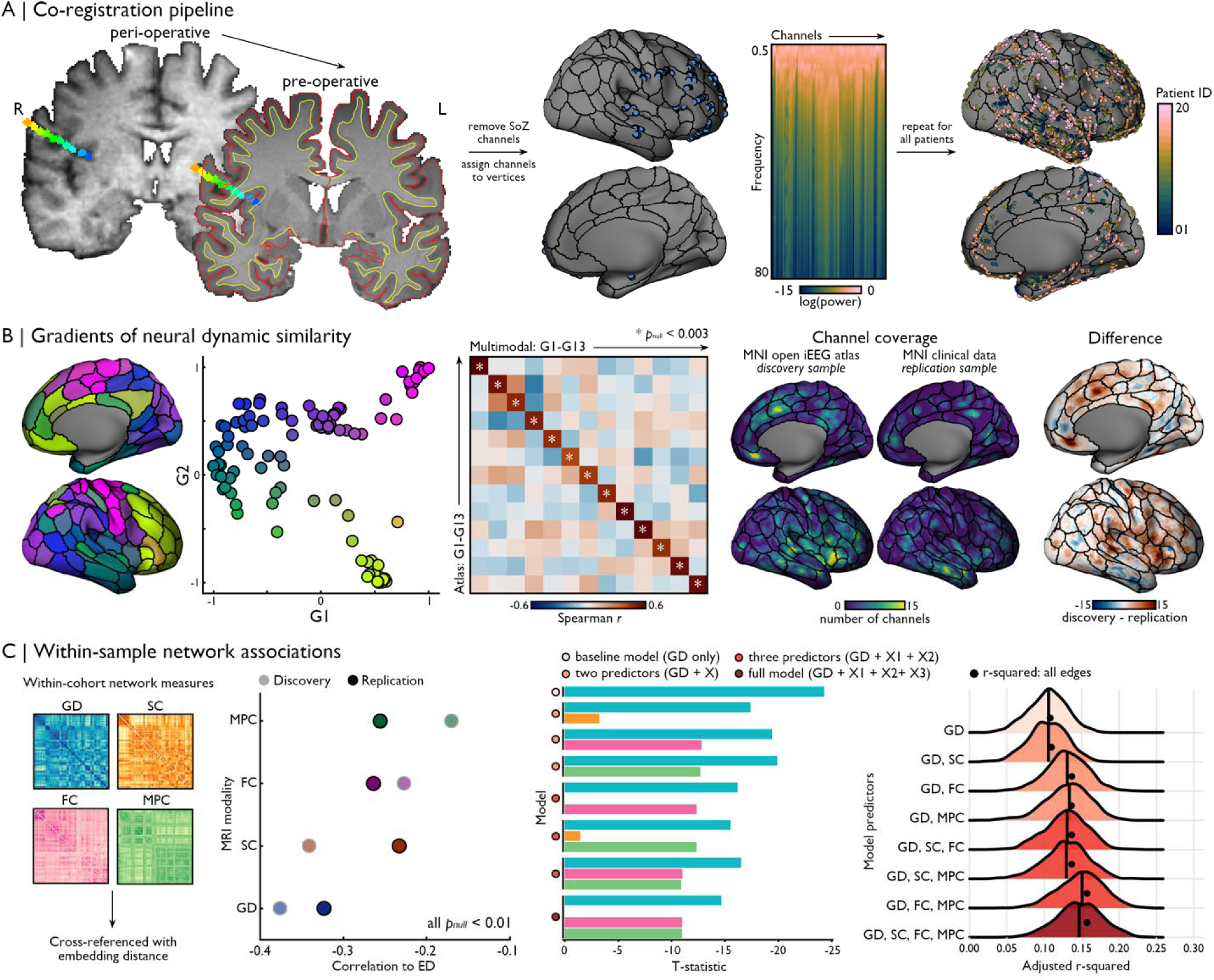
Out-of-sample replication. **(A)** The MNI clinical dataset (n=18) served as a Replication cohort for our gradient analysis pipeline and underwent identical pre-processing as the MNI open iEEG dataset. The coloured markers overlaid on the T1-weighted images indicate the locations of contacts of a single electrode implanted in this patient (deeper contacts in blue, superficial contacts in orange. SoZ: seizure onset zone). **(B)** Gradients of neural dynamic similarity derived from the Replication cohort were all significantly correlated with their counterparts in the Discovery dataset (*p_spin_*<0.05; Bonferroni-corrected *p*<0.003 is indicated by *; one-sided), despite variations in channel coverage across samples. **(C)** We harnessed within-sample measurements of network metrics including, as in our main analyses, GD, SC, FC, and MPC. Bivariate relationships were strongest with GD, and relatively balanced across other network measures. The colour of the dots (correlations to ED) and bars (T-statistic) correspond to the different MRI modalities shown in the left-hand portion of the panel (blue: GD; orange: SC; pink: FC; green: MPC). Coloured dots along the *y*-axis denote model category as shown in Figure 3A. Ridge plots show that variance explained (adjusted r^2^) was stable across models. Each ridge plot shows the distribution of bootstrap results, with their median value indicated by the black line. The black dot marker indicates the true adjusted r^2^ for each model. *Abbreviations*: GD: Geodesic distance; SC: Structural connectivity; FC: Functional connectivity; MPC: Microstructural profile covariance; ED: Embedding distance. Source data are provided as a Source Data file.

Bivariate associations between each MRI modality and embedding distances specific to the Replication cohort again highlighted strongest correlations with GD (r=-0.323; *p_null_*=0.002), and relatively balanced correlations with SC (r=-0.233; *p_null_*<0.001), FC (r=-0.264; *p_null_*<0.001), and MPC (r=-0.256; *p_null_*<0.001) (**Figure 4C**). This stronger contribution of GD was replicated in all multilinear models. The contribution of SC in these models was negligible when either FC or MPC were included as model predictors. Of all modalities, SC was the most dissimilar across cohorts (cross-cohort correlations for each matrix: r_GD_=0.999; r_SC_=0.847; r_FC_=0.960; r_MPC_=0.947). Bivariate correlations between Replication cohort embedding distance and connectome matrices computed from our sample of healthy participants (as used in the Discovery cohort) recovered the overall pattern of findings observed in the Discovery cohort, with descending correlation strength with GD (r=-0.323), SC (r=-0.268), FC (r=-0.250), and MPC (r=-0.245). These findings indicate modality-specific idiosyncrasies potentially sensitive to epilepsy-related connectome alterations when comparing findings across datasets. Leveraging within-sample data furthermore highlighted overall improved prediction of neural dynamics similarity over all edges with increasing model complexity, particularly when including FC and MPC as model predictors (**Figure 4C**). When using iEEG and connectivity data extracted from distinct samples (as done in our Discovery cohort analysis), this effect was only observed in transmodal cortices and when selectively fitting model at the node-level. Additional replication analyses implemented at the individual level also found stronger associations with GD relative to other modalities, followed by SC and balanced correlations with FC and MPC (**Figure S11**). Furthermore, reconstructing the embedding space while iteratively excluding each patient found that embedding distances between nodes related to channel-specific PSD similarity in th held-out patient (**Figure S12**). Together, these findings demonstrate the overall replicability of our main findings across cohort as well as within-patient generalizability of fidings, while additionally identifying nuances in long-range wiring principles of neural dynamics in different analytical designs (between-versus within-subject).

## Discussion

Experimental and modelling studies have produced detailed accounts of the neural hardware implicated in the generation of sustained cortical dynamics. These investigations have highlighted several microscale properties that modulate circuit level inhibition and excitation producing background rhythms (*24, 68, 69*). On the other hand, system-level principles shaping these signals have received less attention, despite their involvement in the coordination and propagation of brain activity (*25*). In the current work, we leveraged the high spatiotemporal precision of iEEG to first map the macroscale topography of human brain dynamics at rest, and then examine its association to different features of neural organization derived from multimodal MRI connectomics (*18*). Our work capitalized on both multicentric open access data sharing initiatives for iEEG and MRI data (*8, 18*), together with a deeply phenotyped sample of patients that had completed iEEG after multimodal MRI. Based on these resources, we could derive consistent gradients of neural dynamics across the neocortex in two independent cohorts, although cohort-specific effects (*i.e.*, relating to iEEG channel coverage) affected replicability of findings. We cross-referenced these gradients with multiscale network information derived from high-definition structural and functional MRI to study the contributions of each modality. Our analyses showed strongest bivariate relationships with geodesic distance, confirming a strong association between local geometry and cortical function (*30, 70*), that likely relates to the former capturing the predominant role of short-range connectivity in shaping cortical function (*64, 71, 72*). Extension of this approach showed that additional network markers, including microstructural similarity and long-range connectivity, improved predictions of neural dynamics, particularly in transmodal regions. Notably, we found gradual shifts in network scale constraints on neural dynamics along the cortical hierarchy, consistent with known connectivity principles. Collectively, our findings demonstrate that the macroscale organization of neural dynamics can be captured by the network architecture of the cortex, and that different network scales can account for regional activity. This work represents an important step in describing the organization of brain function via direct measurements of neural activity, while clarifying how these functional patterns relate to brain wiring.

Combined investigations of brain structure and function offer a unique perspective into brain organization (*73*). These properties are inter-dependent as brain wiring constrains neural activity at local and global scales, while functional, modulatory, and plastic processes shape the brain’s anatomical connections (*74–77*). Mapping the topography of structure-function relationships can, thus, ultimately inform us on the unique properties that enable a given region to support specific aspects of cognition and behaviour (*35*). Extending previous work focusing on local microarchitecture (*20, 21*) and functional measures relying on indirect measurements of neural activity (*78*), the present work comprehensively charted the wiring backbone of inter-regional similarity of intracranially measured neural dynamics across the cortex. Of all queried modalities, we found strongest correlations between inter-regional similarity in neural dynamics and geodesic distance computed along the cortical surface. This finding was replicated using multilinear analyses, as well as when cross-referencing gradients of neural dynamics with geodesic distances in between- and within-cohort designs. Geodesic distance is an intrinsic surface property constraining the appearance and shape of the cortical sheet (*33*). This geometry notably scaffolds travelling waves, a phenomenon under which patterns of neural activity propagate in a coordinated manner across different cortical regions. Travelling waves are thought to be primarily mediated by a horizontal system of cortico-cortical axons (*79–81*). Indeed, short-range cortico-cortical connections, that is with a length up to 3cm, directly follow cortical geometry (*31*). Horizontal collaterals connect neighbouring areas without descending into the white matter, while a U-fibre system of fibres descends into the superficial white matter to connect regions separated by a few centimeters. These short-range connections significantly outnumber long-range fibre tracts descending deep into the white matter, making up over 90% of all cortico-cortical connections (*31*). Given the preponderance of these types of connections, simple wiring rules based on anatomical neighborhoods can approximate overall cortical wiring (*82–85*) and functional connectivity (*64, 71, 72*). In line with recent evidence demonstrating the significant contribution of cortical geometry in constraining macroscale function (*30*), our findings highlight a stronger relationship between inter-regional geodesic distance and neural dynamics similarity over long-range connectivity metrics.

Prior work on the topography of functional network architectures has demonstrated a shifting balance of short- to long-range connectivity profiles along the cortical hierarchy. Specifically, the activity of heteromodal cortices – at high hierarchical distance from primary sensory regions – is strongly coupled with the activity of spatially distributed regions (*86–89*). This higher functional connectivity distance of association cortices is reflected in the connectivity architecture of *hub* regions, which form a heavily interconnected wiring backbone that cannot solely be explained by rules of anatomical proximity (*83, 90*). Following this line of work, we cross-referenced the neural dynamic embedding space with modalities sensitive to macroscale structural and functional network architectures to quantify the contributions of these metrics in addition to those afforded by geodesic distance (*54*). We specifically implemented multilinear models to quantify the relationship between neural dynamics similarity and network properties at a regional scale. We found that models including microstructural profile similarity selectively improved model accuracy in transmodal regions, and that these results could be further augmented by the inclusion of either structural or functional connectivity metrics. While some redundancy exists across these networks measures, our analysis suggests that the similarity of local microcircuits properties reflected by microstructural profile covariance may be a key contributor to patterns of neural dynamics (*20*). Coupled with regional connectivity modelled by geodesic distance, microstructural similarity may thus reflect the influence of local microstructure on ongoing neural activity.

Although the present work emphasized multiscale network-level associations, intra-regional properties are also likely contributors to ongoing neural dynamics. In our findings, the two gradients explaining the most variance in neural dynamic similarity differentiated systems along sensory-motor (G1) and unimodal-transmodal (G2) axes. These patterns are reminiscent of those found in other structural and functional data modalities (*91*) and recapitulate established models of information processing linking extrapersonal (sensory input and motor output) and internal spaces (transmodal limbic and paralimbic structures) (*61*). Capturing higher-frequency signals in fronto-motor regions and high delta to low alpha peaks of temporo-occipital areas, G1 may reflect signal properties segregating perception and action systems. Conversely, the cortical processing hierarchy running from unimodal to transmodal regions has repeatedly emerged from the analysis of gradients of intrinsic functional connectivity derived from rs-fMRI (*56*). This axis was recovered here in G2, although our approach aggregating signals from multiple different patients did not quantify functional connectivity. The transmodal anchor of G2 was found to exhibit stronger low-frequency activity, particularly in the delta range. Consistent with this finding is the identification of longer neural timescales of information processing in these regions, as quantified by signal autocorrelation analysis (*21, 89, 92–96*), thought to be key for processes of information maintenance over longer periods of time (*97*).

These complementary axes of neural dynamic similarity may be supported by overlapping principles of cortical microstructural differentiation. Regarding G1, histological studies have shown that sensory and motor cortices are characterised by opposing progressive shifts in cortical and laminar thickness (*98–101*). Indeed, layer IV increasingly thickens when going from primary motor to anterior frontal regions, while thickness of layers III, V, and VI as well as the overall thickness of the cortex gradually decrease in this direction. In sensory hierarchies involving visual, somatosensory, and auditory systems, the opposite trend is observed. These changes are thought to reflect underlying shifts in laminar cytoarchitecture, as the size of cells and dendritic arbours, as well as the number and density of synapses tend to increase with cortical thickness (*102–104*). Together, these features contribute to a larger neuropil volume and concurrent cortical thickening (*98*), suggesting a local microarchitectural basis of signal variations captured by G1, which we could confirm through this gradient’s specific association to cytoarchitectonic classes (*58*). Conversely, the unimodal-transmodal gradient captured by G2 seems to co-vary with large-scale shifts in cortical laminar elaboration (*35*). For instance, the length of neural timescales across the cortex is inversely correlated with a putative measure of intracortical myelination (*21*), reflecting variations in local microarchitecture but also co-occurring changes in large-scale connectivity patterns (*93, 105–107*). Together, these findings may suggest a changing balance of local and network-level properties in constraining neural function that can be tracked by local microarchitecture.

Cross-referencing iEEG data collected in patients with MRI measures acquired in healthy participants allowed us to maximize spatial coverage and sample sizes in our Discovery cohort analyses. This approach is in line with the widespread use of data contextualization techniques in neuroscience, which measures and integrates distinct maps of human brain organization (*108*). However, this remains a limitation of the present work, as it is unclear to what extent findings from these distinct cohorts generalize to overall principles of brain organization. Indeed, translating our approach at an individual-level revealed considerable inter-individual variability in the magnitude of multimodal association. However, given the technical limitations of iEEG, it is difficult to disentangle the relative contributions of individual differences in brain organization versus spatial sampling effects in explaining this variability. We were generally able to replicate our main results in a deeply phenotyped independent cohort in which both invasive neurophysiology as well as multimodal imaging data were available in the same patients. Some differences were nevertheless observed across cohorts. First, bivariate correlations in the Replication cohort showed a stronger relationship between neural dynamic similarity and microstructural similarity, and a lower correlation with structural connectivity relative to the Discovery cohort. In our multilinear approach, the contribution of structural connectivity was generally minimal in the Replication cohort. This may be due to a multitude of factors, including more precise co-registration and cross-modal alignment in the Replication cohort, differences in channel coverage that may reflect different clinical features of both Discovery and Replication cohorts (e.g., the higher coverage in temporal, relative to frontal, insular, and parietal channels, in the Replication cohort; slight overrepresentation of women in the Replication cohort), as well as epilepsy-related changes in SC that could only be detected in a within-patient design. More broadly, differences in socio-demographic profiles (*e.g.*, slight overrepresentation of women in the Replication cohort vs. a more balanced sex ratio in the Discovery cohort) may have influenced resulting iEEG gradients as well as MRI-derived measures. We could however recover overall patterns of correlations between connectome modalities and the neural dynamic embedding space by using connectome matrices from our normative cohort and gradients computed from the Replication cohort. These findings highlight some of the limitations of cross-dataset contextualization, which remains a correlational approach limited in its potential to deliver causal insights into individual-level mechanisms of brain organization. Biophysical modelling techniques may be suitable for these investigations, given their ability to identify local and network parameters regulating brain function (*92, 109, 110*) and to study how perturbations of these parameters can skew functional connectome organization (*111*). Of interest, recent modelling studies have emphasized that relatively rare long-range connections play a key role in optimizing information flow across brain networks, alongside distance-based principles anchored in cortical geometry (*53, 112*).

Nonetheless, investigations of human brain organization using iEEG are limited by the technique’s unique challenges, including its restricted accessibility, exclusive use in clinical settings, as well as sparse and patient-specific sampling (*113*). It also remains an open question to what extent focal epileptic activity in one brain region might affect activity in the rest the brain and skew the generalizability of iEEG findings to healthy brain function. Recent data sharing initiatives, such as the resource used here (*18*) and others (*8, 114–117*), have been instrumental in addressing these limitations. Although iEEG data is collected in a clinical context, pooling carefully curated single-patient data significantly boosts spatial coverage, thus enabling explorations of brain cartography and facilitating the identification of overarching principles of neural organization. These innovations add on to the recognized advantages of iEEG regarding its high spatiotemporal precision to study human brain function, as well as its specificity to neural signals over methods offering full brain coverage such as fMRI (*113*). Studies focusing on group-level inference, such as the present work, thus offer new perspectives for human brain mapping to which other modalities may be blind. Group-level paradigms are a valuable step towards more targeted individual-level inquiries able to clarify populational variations. For example, by testing the consistency of these principles across individuals, extreme deviations from the population norm can be identified and related to clinical presentations (*3*). Future studies based on even larger datasets or providing a high proportion of bilaterally implanted cases may furthermore help to elucidate principles of asymmetry in neural dynamic across the two hemispheres.

Despite these caveats, the present work offers a multiscale view of network constraints on human neocortical dynamics. We particularly highlight the key role of open datasets in facilitating these investigations, notably allowing us to sidestep a significant limitation of iEEG data by boosting spatial coverage of the cortical sheet via a multisite data sharing initiative (*18*). This resource helped us address an important gap in the literature concerning the basis of cortical neural dynamics. By extending investigations to iEEG and MRI, our findings solidify evidence positioning unimodal-transmodal differentiation as an overarching principle of brain organization recapitulated in cortical wiring and dynamics. Importantly, this work provides a comprehensive, cortex-wide description of these dynamics and their relationship to multiscale network architectures.

## Methods

The present work relies on open-source datasets. The studies under which these datasets were collected complied with the Research Ethics board of McGill University and the Montreal Neurological Institute and Hospital. Written and informed consent were obtained from all participants to take part in the research MRI protocol, and they received monetary compensation for their participation in MRI studies. As iEEG data was acquired as part of each patient’s clinical workup for intractable epilepsy, patients were not compensated for this aspect of data collection.

### Dataset overview

*a) Atlas of the human intracranial electroencephalogram (Discovery cohort).* The MNI open iEEG atlas is a multicentre initiative providing openly available human intracranial recordings acquired during different states of vigilance (*8, 18*). This dataset includes iEEG recordings from a sample of 106 patients (52 women; mean ± standard deviation (SD) age = 33.1±10.8 years) with intractable epilepsy, totalling over 1,700 data points taken from non-epileptic channels mapped to a common stereotactic space. The present study focused on segments recorded during resting wakefulness with eyes closed. All data was collected during a standard clinical protocol implemented in all participating sites.

*b) MNI clinical dataset (Replication cohort).* We studied a sample of 18 patients (11 women; mean ± SD age = 33.89±9.02 years; mean ± SD age at seizure onset 13.28±7.47) treated at the Montreal Neurological Institute and Hospital for intractable epilepsy. Patients underwent multimodal MRI at a field strength of 3 Tesla as part of a research protocol prior to iEEG investigations completed as part of their clinical workup. Patients showed heterogeneous suspected locations (frontal: n=3; temporal: n=6; parietal: n=4; occipital: n=1; insula: n=4) and laterality (right: n=11; left: n=6; bilateral: n=1) of the epileptogenic focus. An epileptologist (TA) reviewed clinical and iEEG data and labelled channels in the seizure onset zone, which were excluded from our analysis. Nine patients showed suspected focal cortical dysplasia (FCD; histopathological confirmation of FCD type IIa in six patients; FCD type I in one patient) and one patient presented with a ganglioglioma, while the others were deemed MRI-negative with no available surgical tissue for histopathological analysis. 13/18 patients underwent surgery to remove the suspected epileptic focus (surgical resection or radiofrequency thermocoagulation), with eight patients achieving seizure freedom after surgery (Engel Ia or Ib).

### Electrophysiological data pre-processing and gradient analysis

*a) iEEG data pre-processing.* All data segments from bipolar channels provided in the MNI open iEEG atlas underwent identical pre-processing (*18*). In brief, these steps included the application of a band-pass filter at 0.5 to 80Hz, downsampling to 200Hz, application of an adaptive filter to reduce powerline interference (50 or 60Hz depending on site) and demeaning each data segment. When no continuous and artefact-free 60-second segments were found for a given channel, multiple discontinuous segments from the channel were concatenated with a 2-second zero-padded buffer between segments. Following the procedure established in this dataset, we used Welch’s method to compute each channel’s PSD from 2-second blocks with 1-second overlap, weighted by a Hamming window. Each channel’s PSD was normalized so the total power was equal to one. As for the MNI clinical dataset, two board-certified epileptologists (BF, DM) selected data segments (60 to 120 second duration) free of ictal events and artefacts from periods of resting wakefulness with eyes closed. Like the MNI open iEEG atlas, selected epochs could be continuous or discontinuous, in which case data segments were concatenated with a 2-second zero-padded buffer. All recordings were acquired at a sampling rate of 2000Hz but were downsampled to 200Hz to match the pre-processing applied to the MNI open iEEG atlas. This cut-off allowed us to cover commonly studied frequency bands (from delta to gamma range) while eliminating high frequency noise. However, this also filters potentially high frequency content such as physiological ripples and fast ripple activity (*118*).

*b) Anatomical localization and co-registration of channels.* For the MNI open iEEG atlas, all channel coordinates were provided in a common stereotactic space (ICBM 2009a symmetric) alongside symmetric left and right cortical surface meshes (119k vertices per hemisphere). As this surface is symmetric, channels located in the left hemisphere were flipped to the right hemisphere to improve spatial coverage. As a result, all downstream analyses, including the computation of gradients of neural dynamics and correlations to MRI metrics, are implemented in a single hemisphere. Each channel was attributed to the closest vertex on the cortical surface determined by the shortest Euclidean distance between each vertex and the channel’s position in volumetric space. To account for effects of volume conduction, channel PSDs were extrapolated to surrounding vertices within a 5mm radius (*channel region*). Distances were measured using the exact geodesic algorithm (*119*) implemented in the Pygeodesic package (v0.1.7). Channel regions underwent surface-based registration to the fsLR surface template (∼32k vertices) using nearest-neighbour interpolation.

For each patient in the MNI clinical dataset, peri-implantation scans were co-registered to pre-surgical T1w images pre-processed using micapipe (v0.2.3) (*120*). We applied a label-based affine registration (*121, 122*) to align peri-implantation T1w (9/18 patients) and T2w (2/18 patients) to pre-surgical native space. When only computerized tomography (CT) was available for peri-implantation imaging (6/18 patients), scans were directly aligned to pre-surgical space without prior tissue-type segmentation. We applied the resulting transform to each patient’s electrode contact coordinates, bringing them to patient-specific native pre-surgical space. Retaining only contacts overlapping with neocortical grey matter, channel coordinates were defined as the midpoint between the two contacts making up the channel. Neocortical surface segmentations were generated from pre-surgical T1w scans using FreeSurfer 6.0 (*123–125*), and were resampled to retain the geometry of the fsLR surface template (∼32k vertices). Channels were attributed to the closest vertex on the neocortical surface mesh as measured with Euclidean distance (*channel vertex*). As with the MNI open iEEG atlas, channel PSDs were extrapolated to surrounding vertices within a 5mm radius (*channel region*) as defined by the exact geodesic algorithm (*119*) implemented in the Pygeodesic package (v0.1.7). Channels involved in seizure onset (*i.e.,* seizure onset zone) were identified by an experienced epileptologist (TA) and excluded from further analyses.

*c) Generating gradients of neural dynamics.* Channel PSDs were log-transformed and mapped to the right hemisphere labels of the Schaefer-200 parcellation (*57*), resulting in 100 parcel-wise PSDs. This parcellation resolution was selected as it provided complete sampling of the right hemisphere. To map channel-level PSDs to this parcellation, we first identified all channel regions overlapping with a given parcel. For each frequency (0.5-80Hz, in increments of 0.5), the amplitude of the PSD for all channels covering the parcel was averaged. More specifically, we implemented a weighted average where amplitudes were weighted according to the extent of the channel region’s coverage of the parcel (the number of vertices that channel contributed to the parcel), resulting in a parcel-level PSD. Parcel-wise PSDs were cross-correlated while controlling for the average PSD computed across all channels and underwent Fisher R-to-Z transformation. We used the BrainSpace toolbox (*126*) to convert the Z-transformed correlation matrix to a normalized angle affinity matrix, and applied diffusion map embedding to generate eigenvectors or *gradients* of neural rhythm similarity across the cortex using default sparsity (keeping only the top 10% of weights) and diffusion (α=0.5) parameters. Of note, systematic correlations spanning the gradient hyperparameter space found the resulting gradients to be generally robust to parameter selection in both Discovery and Replication cohorts (**Figure S7**). The same pipeline was applied in the Replication sample, which was additionally aligned to the embedding space of the Discovery dataset using Procrustes alignment. We compared two complementary alignment strategies. First, using singular value decomposition, we computed a transformation matrix enabling translation, rotation, and scaling of the Replication cohort gradients to optimize their similarity to the Discovery cohort gradients. The results of this approach are presented in **Figure 4**. We also applied a more conservative approach where gradients from the Replication cohort were simply reordered to minimize their distance to gradients computed in the Discovery cohort (**Figure S10**). Consistency of gradient topographies across datasets was quantified using Spearman rank correlations and statistical significance was determined using spin permutation tests.

### MRI data acquisition

a) *Participants.* Data were collected in a sample of 50 healthy volunteers (23 women; age mean ± SD = 30.28±5.74 years) denying any history of neurological and psychiatric illness. The Ethics Committee of the Montreal Neurological Institute and Hospital approved the study. Raw and pre-processed connectomes data are openly available as part of the MICA-MICs dataset (*55*) (<u>osf.io/j532r/</u>, note that updated outputs generated with micapipe v0.2.3 (*120*) were used in the present study). Patients included in the MNI clinical dataset (Replication cohort) underwent the same multimodal imaging protocol as healthy volunteers.

b) *MRI acquisition*. Scans were acquired at the McConnell Brain Imaging Centre of the Montreal Neurological Institute and Hospital on a 3T Siemens Magnetom Prisma-Fit equipped with a 64-channel head coil. We acquired two T1-weighted (T1w) scans with identical parameters using a 3D magnetization-prepared rapid gradient-echo sequence (MPRAGE; 0.8 mm isovoxels, TR=2300 ms, TE=3.14 ms, TI=900 ms, flip angle=9°, FOV=256×256 mm^2^). T1w scans were inspected to ensure minimal head motion before undergoing further processing. Quantitative T1 (qT1) relaxometry data were acquired using a 3D-MP2RAGE sequence (0.8□mm isotropic voxels, 240 sagittal slices, TR□=□5000□ms, TE□=□2.9□ms, TI 1□=□940□ms, T1 2□=□2830□ms, flip angle 1□=□4°, flip angle 2□=□5°, iPAT□=□3, bandwidth□=□270□Hz/px, echo spacing□=□7.2□ms, partial Fourier□=□6/8). We combined two inversion images for qT1 mapping to minimise sensitivity to B1 inhomogeneities and optimize reliability. (*127, 128*) A 2D spin-echo echo-planar imaging sequence with multi-band acceleration was used to obtain DWI data, consisting of three shells with b-values 300, 700, and 2000s/mm^2^ and 10, 40, and 90 diffusion weighting directions per shell, respectively (1.6mm isotropic voxels, TR=3500ms, TE=64.40ms, flip angle=90°, refocusing flip angle=180°, FOV=224×224 mm^2^, slice thickness=1.6mm, multi-band factor=3, echo spacing=0.76ms). b0 images acquired in reverse phase encoding direction were used for distortion correction. One 7□min resting-state function MRI (rs-fMRI) scan was acquired using multiband accelerated 2D-BOLD echo-planar imaging (3□mm isotropic voxels, TR□=□600□ms, TE□=□30□ms, flip angle□=□52°, FOV□=□240□×□240□mm^2^, slice thickness□=□3□mm, mb factor□=□6, echo spacing□=□0.54□ms). Participants were instructed to keep their eyes open, look at a fixation cross, and not fall asleep. Distortion correction of rs-fMRI data was performed using two spin-echo images with reverse phase encoding (3□mm isotropic voxels, TR□=□4029□ms, TE□=□48□ms, flip angle□=□90°, FOV□=□240□×□240□mm^2^, slice thickness□=□3□mm, echo spacing□=□0.54□ms, phase encoding□=□AP/PA, bandwidth□=□2084□Hz/Px).

c) *<u>Replication 7T MRI dataset</u>*. Data were collected in a sample of 10 healthy volunteers (6 women; 26.60 ± 4.60 years) denying any history of neurological and psychiatric illness. Raw and pre-processed connectomes data are openly available as part of the MICA-PNI dataset. Only the first available study session for each participant was analysed as part of this study. Detailed acquisition and pre-processing parameters are available in the MICA-PNI descriptor article (*65*).

### MRI data processing

Imaging data were processed with micapipe (v0.2.3; <u>micapipe.readthedocs.io/</u>), an openly available pipeline for multimodal MRI processing, data fusion, and multiscale connectome generation (*120*). This pipeline is notably adapted to sample features in modality-specific spaces, therefore limiting data blurring that can occur with co-registration to a standard template space.

*a) T1w pre-processing and geodesic distance measurements*. Structural processing was carried out using several software packages, including tools from AFNI, FSL, and ANTs (*129*). Each T1w scan was deobliqued and reoriented to standard neuroscience orientation (LPI: left to right, posterior to anterior, and inferior to superior). Both scans were then linearly co-registered and averaged, automatically corrected for intensity nonuniformity (*130*), and intensity normalized. Cortical surface segmentations were generated from native T1w scans using FreeSurfer 6.0 (*123–125*). We computed individual GD matrices along each participant’s native cortical midsurface using workbench tools (*131, 132*), specifically the *-surface-geodesic-distance* function. First, a centroid vertex was defined for each cortical parcel by identifying the vertex with the shortest summed Euclidean distance from all other vertices within its assigned parcel. The GD between centroid vertices and all other vertices on the native midsurface mesh was computed using Dijkstra’s algorithm. Notably, this implementation computes distances not only across vertices sharing a direct connection, but also across pairs of triangles which share an edge to mitigate the impact of mesh configuration on calculated distances. Vertex-wise GD values were then averaged within parcels. Of note, longer distances measurements may be obtained here due to our use of high-resolution meshes, compared to techniques using surfaces resampled according to a lower-resolution template’s geometry.

*b) DWI pre-processing and tractography-derived SC*. DWI data were pre-processed using MRtrix (*133, 134*). DWI data were denoised (*135, 136*), underwent b0 intensity normalization (*130*), and were corrected for susceptibility distortion, head motion, and eddy currents using a reverse phase encoding from two b=0s/mm^2^ volumes. Required anatomical features for tractography processing (*e.g.,* tissue type segmentations, parcellations) were non-linearly co-registered to native DWI space using the deformable SyN approach implemented in ANTs (*137*). Diffusion processing was performed in native DWI space. Structural connectomes were generated with MRtrix from pre-processed DWI data (*133, 134*). We performed anatomically-constrained tractography using tissue types (cortical and subcortical grey matter, white matter, cerebrospinal fluid) segmented from each participant’s pre-processed T1w images registered to native DWI space (*138*). We estimated multi-shell and multi-tissue response functions (*139*) and performed constrained spherical-deconvolution and intensity normalization (*140*). We generated a tractogram with 40M streamlines (maximum tract length=250; fractional anisotropy cutoff=0.06). We applied spherical deconvolution informed filtering of tractograms (SIFT2) to reconstruct whole brain streamlines weighted by cross-sectional multipliers (*141*). The reconstructed cross-section streamlines were mapped to the Schaefer-200 parcellation, which was also warped to DWI space using the previously generated transforms (*137*). The connection weights between nodes were defined as the weighted streamline count. Group-level structural connectivity (SC) was computed using distance-dependent thresholding to preserve the distribution of within- and between-hemisphere connection lengths found in individual subjects (*142*). However, SC weights of the resulting matrix remain highly skewed, with few very strong connections and many weak or zero connections. As this property of the matrix violates assumptions of normality, which is problematic for the statistical tests applied in the present work, connection weights in the resulting group-level were log-transformed to reduce connectivity strength variance.

*c) Resting-state functional connectivity (FC)*. After discarding the first five volumes of each subject’s rsfMRI run to ensure magnetic field saturation, images underwent reorientation, as well as motion and distortion correction. Motion correction was performed by registering all timepoint volumes to the mean volume, while distortion correction leveraged main phase and reverse phase field maps. Nuisance variable signal was removed using an ICA-FIX (*143*) classifier and by regressing timepoints with high motion detected from motion outlier outputs provided by FSL. Volumetric timeseries were averaged, and the mean volume was registered to each subject’s T1w image using a label-based affine registration (*121, 122*). We applied the resulting transform to previously computed surface files to sample volumetric timeseries using trilinear interpolation. Timeseries were averaged within nodes defined by the Schaefer-200 parcellation and were cross-correlated to generate individual-specific functional connectomes. Correlation values (Pearson’s *r*) subsequently underwent Fisher-R-to-Z transformations. Group-level functional connectivity (FC) was computed by averaging matrices across participants.

*d) qT1 pre-processing and microstructural profile covariance (MPC)*. A series of 14 equivolumetric surfaces were constructed for each participant between pial and white matter boundaries. We aligned qT1 and native FreeSurfer space T1w volumes using a label-based affine registration (*121, 122*). We applied the resulting transform to previously computed equivolumetric surface files to sample image intensities in native qT1 space. This resulted in vertexwise microstructural intensity profiles, which were used to calculate individual microstructural profile similarity matrices (*35, 37*). Intensity profiles were averaged within parcels, cross-correlated across using partial correlations controlling for the average cortex-wide intensity profile, and log-transformed (*37, 48*). Resulting matrices thus represented participant-specific similarity matrices in myelin proxies across the cortex. Matrices were subsequently averaged for group-level analyses.

### Multiscale connectomics of neural dynamics

*a) Bivariate associations between neural dynamics and multiscale network metrics*. The similarity of neural dynamics was quantified by computing the Euclidian distance between all node pairs in the gradient space. For this purpose, we retained gradients explaining a cumulative sum of 50% of the variance (G1-G14; note that results are displayed in a 2D representation composed of G1 and G2). Node proximity in the embedding space was cross-referenced with measures of *(i)* cortical geometry indexed by GD, *(ii)* tractography-based SC, *(iii)* resting-state FC and *(iv)* microstructural similarity (MPC) by correlating the upper triangular edges of each matrix. For each modality, statistical significance was determined by comparing empirically observed correlations to those obtained with surrogate matrices in which each row had been shuffled using spatial autocorrelation-preserving null models (*63, 64*). Consistency of cross-modality associations were also assessed by comparing embedding distance distributions across thresholded network matrices preserving the 5% to 95% strongest edges (in increments of 5%). We also assessed node-level bivariate associations by repeating correlations independently for each matrix row, and cross-referencing resulting correlation coefficients with those obtained spatial null models (*63, 64*).

*b) Multiple linear regression models*. Multiple linear regression models were implemented with distinct sets of modality predictors (*i.e.*, a baseline model including GD only; partial models including GD and one other modality, or two other modalities from: SC, FC, or MPC; a full model including all four modalities). Models were additionally implemented using either all edges of the upper triangular part of each matrix or on a region-specific basis. Goodness-of-fit of each model was assessed using adjusted r-squared. In models implemented over all edges, we compared the importance of each feature from its t-statistic, *i.e.* each feature’s coefficient divided by its standard error. We assessed the generalizability of model parameters by bootstrapping, specifically subsampling 90% of edges to estimate model parameters and testing the goodness-of-fit over the remaining 10% of edges. Region-specific models were also implemented to quantify node-level gains in adjusted r-squared, which were spatially correlated with the principal gradient of resting-state functional connectivity derived from an independent sample as provided in the BrainStat toolbox (*56, 66*).

## Data availability

Source data are provided with this paper. The authors acknowledge the use of several open datasets in making the present work possible. The MNI open iEEG atlas data are available at https://ieegatlas.loris.ca/ provided a user account creation (free), the MICA-MICs data used in this study are openly available (https://osf.io/j532r/), and the MICA-PNI data used in this study are also openly available (https://osf.io/mhq3f/).

## Code availability

Code to reproduce the main analyses of this paper, alongside fully processed data dependencies, are available on Github (https://github.com/royj23/ieeg_gradients). This code relies on several open software, mainly micapipe (https://github.com/MICA-MNI/micapipe) for 3T and 7T MRI data pre-processing as well as BrainSpace (https://github.com/MICA-MNI/BrainSpace) for the computation of cortical gradients. Visualization of cortical surface data rely on an openly available package (https://github.com/StuartJO/plotSurfaceROIBoundary).

## Acknowledgements and funding

J.R. is supported by a fellowship from the Canadian Institutes of Health Research (CIHR). C.P. is supported by the Deutsche Forschungsgemeinschaft (DFG Emmy Noether Programme-524408221). E.S. acknowledges funding from the Fonds de Recherche du Québec – Santé (FRQS) Doctoral Training Scholarship. B.F. acknowledges support from CIHR (PJT-175056) and a salary award (Chercheur-boursier clinicien Senior) from the FRQS. B.C.B. acknowledges research support from the National Science and Engineering Research Council of Canada (NSERC Discovery-1304413), CIHR (FDN-154298, PJT-174995, PJT-191853), SickKids Foundation (NI17–039), the Helmholtz International BigBrain Analytics and Learning Laboratory (HIBALL), Healthy Brains and Healthy Lives, BrainCanada, and the Tier-2 Canada Research Chairs program. We additionally thank all patients and healthy volunteers who gave their time and contributed to these datasets.

## Author contributions statement

Conceptualization: J.R., B.C.B; Methodology: J.R., B.C.B.; Patient outreach and recruitment: B.F., R.P., J.H.; Data collection: H.A., A.N., E.S., B.F.; Data curation: J.R., C.P., T.A., C.A., D.M.; Data processing: J.R., C.P, R.R.C., E.S.; Formal analysis: J.R.; Writing - Original Draft: J.R., B.C.B.; Writing - Review & Editing: J.R., C.P, R.L., J.S., B.F., B.C.B; Visualization: J.R.; Supervision: B.C.B.

