## Supplementary figures for "Human cortical dynamics reflect graded contributions of local geometry and network topography"

**Short title**

Gradients of neural dynamics

**Corresponding authors**

Jessica Royer, Psy.D., Ph.D.

Boris C. Bernhardt, Ph.D.

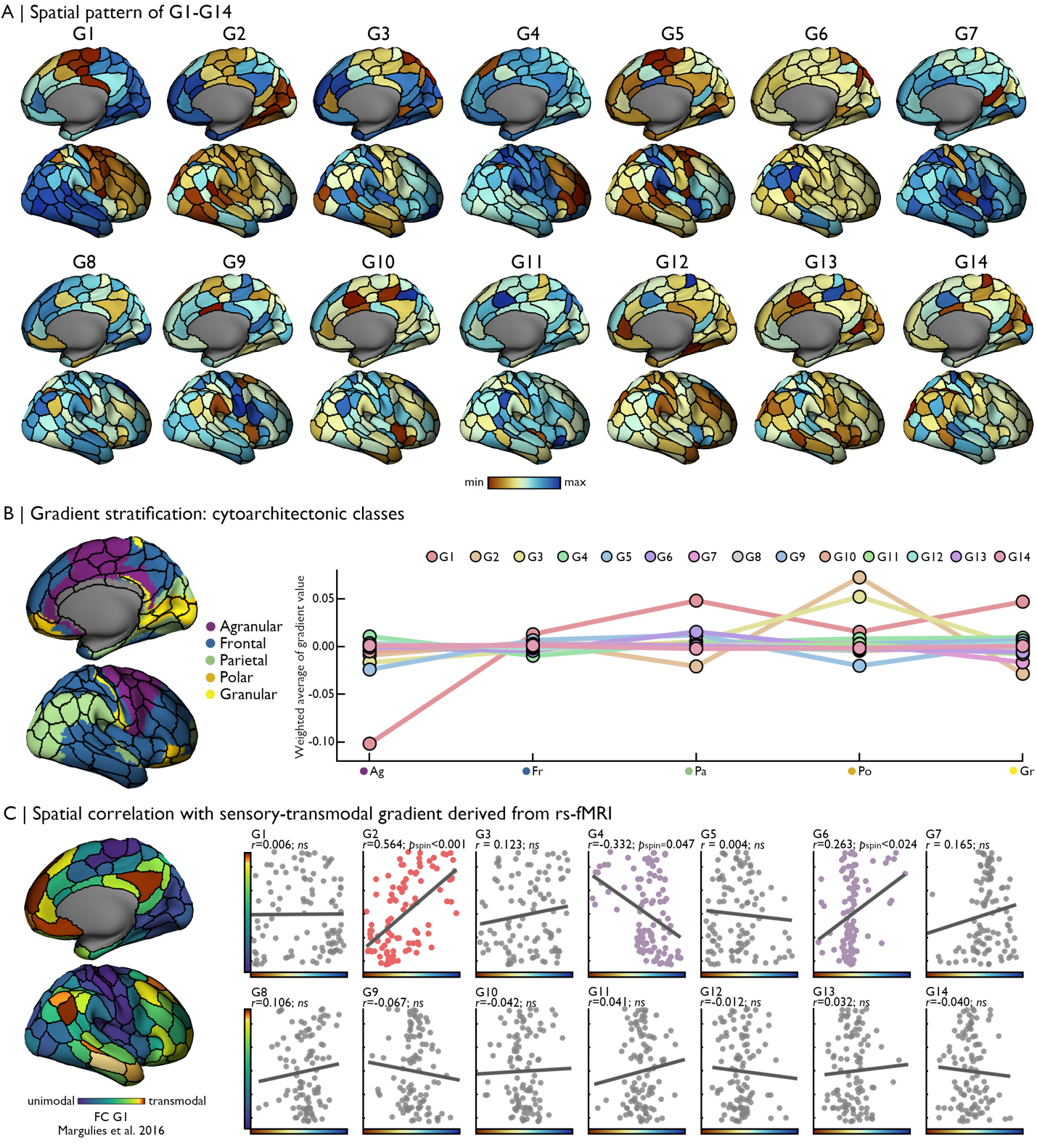

**Figure S1. Gradient topographies. (A)** G1-G14 cumulated to 50% of explained variance in PSD similarity across the cortex and were retained in further analyses. **(B)** For each gradient, we computed the average gradient values of parcels in each class of the Von Economo and Koskinas atlas. Values were weighted by the proportion of vertices in each parcel covered by each cortical type. A simple linear regression fit over the label of each class, reflecting its level of laminar differentiation (1-low; to 5-high), and their corresponding weighted average gradient value show a higher, positive slope for G1 (slope = 0.03). Indeed, G1 most strongly differentiated agranular (purple) from parietal (moss) and granular (yellow) classes. However, slopes for G2-G14 were of much lower magnitude (slope range: -0.002 to 0.009). **(C)** Of all analysed gradients, G2 was most strongly correlated with the unimodal-transmodal axis of cortical organization (pink; Spearman *r*=0.564, *p*_spin_<0.001). Trend-level associations found with G4 and G6 (lavender) did not survive Bonferroni correction. Source data are provided as a Source Data file.

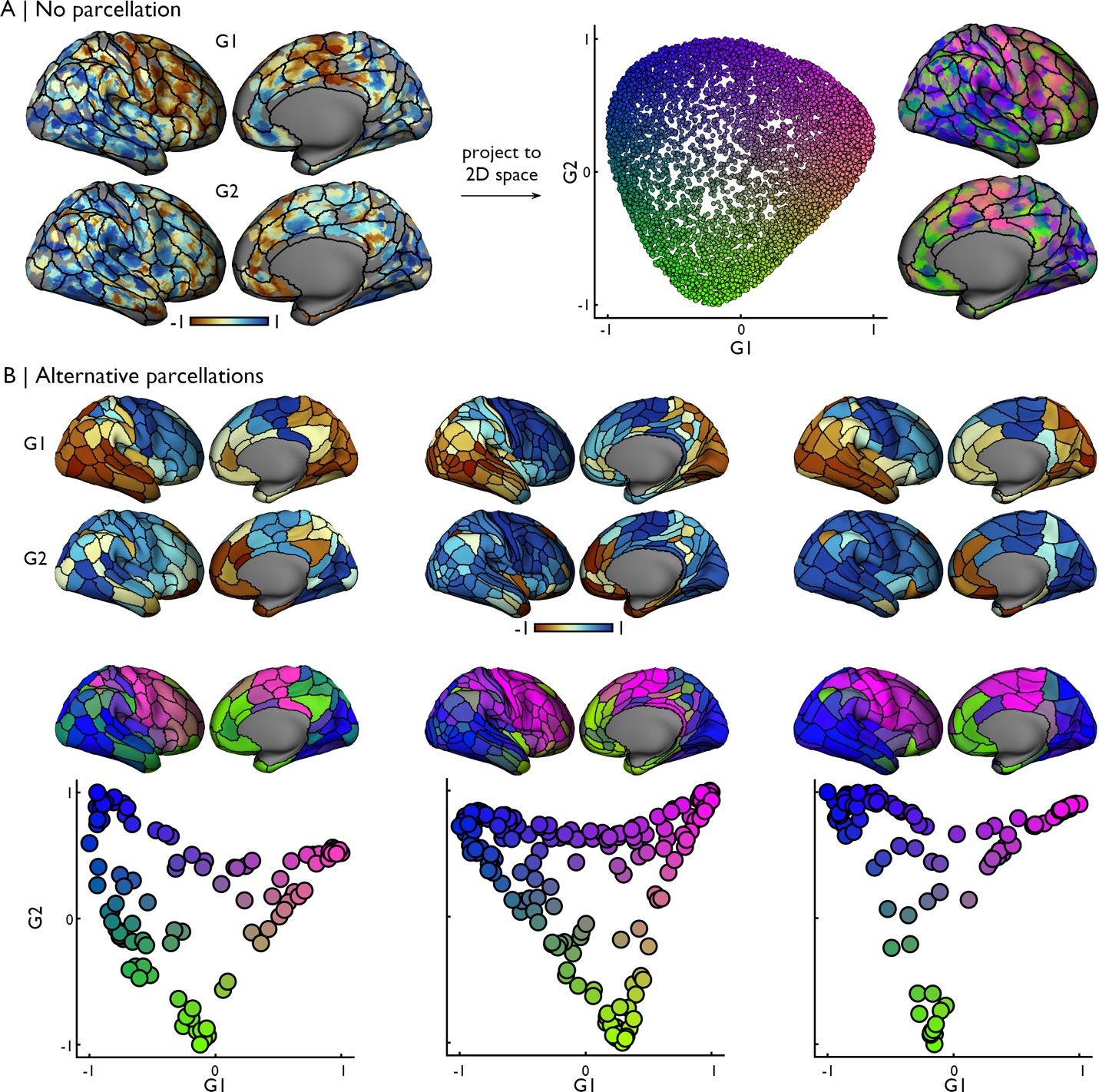

**Figure S2. Consistency across parcellations schemes. (A)** Gradients 1 and 2 when generated from unparcellated data, i.e. “channel regions”. **(B)** Gradients 1 and 2 when generated from different parcellations applied to the PSD data: Schaefer-200 (as in the main manuscript; selected as it provided complete iEEG sampling of the right hemisphere), Glasser-360, and a random 200-node subdivision of the aparc parcellation. Source data are provided as a Source Data file.

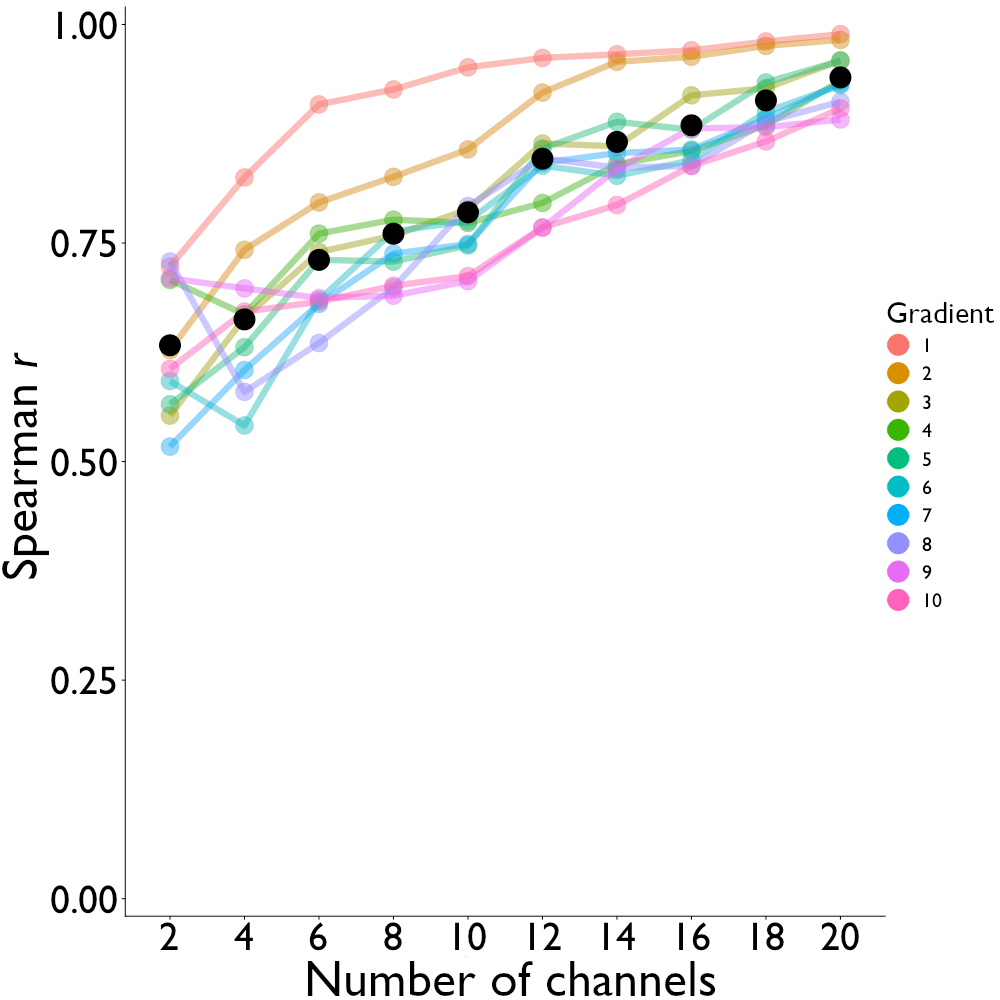

**Figure S3. Replicability across channel subsets.** We repeated our gradient analysis while varying the number of channels included in each parcel prior to averaging PSDs across channels in each parcel. We retained the specified number of channels with the highest vertex coverage in each parcel. Coloured lines indicate Spearman r between replication analysis and original findings in the main manuscript for each gradient at a given channel number. Black dot indicated mean Spearman r across gradients for a given channel number. Source data are provided as a Source Data file.

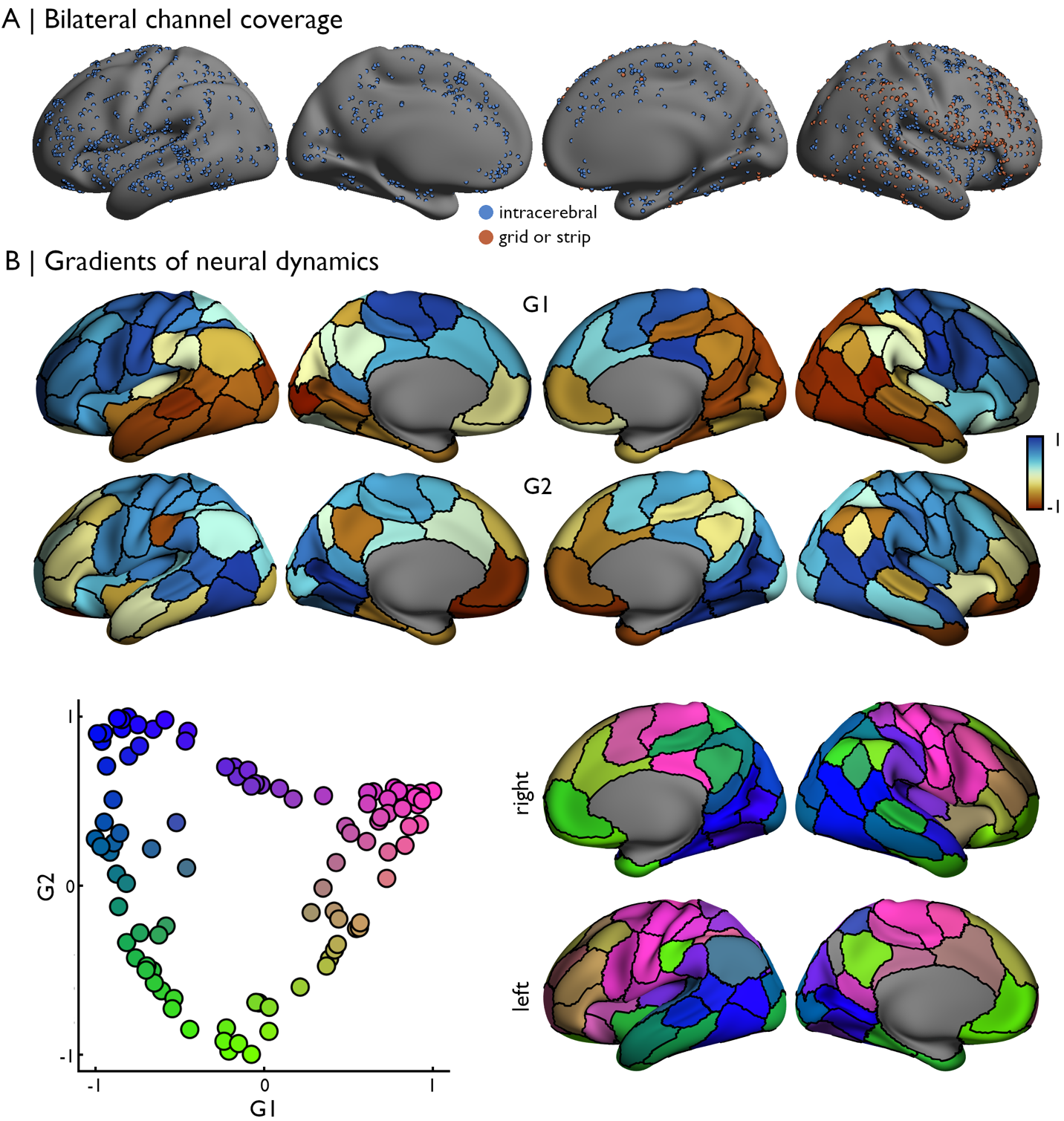

**Figure S4. Consistency of findings when preserving bilateral distribution of channels. (A)** Channel coordinates from the MNI open iEEG atlas were registered to each hemisphere of the fs-LR-32k surface template. Overall, the right hemisphere was move extensively covered by iEEG channels. **(B)** Repeating the same analytical procedure as in the main manuscript, we mapped channel PSDs to a 100-node parcellation (Schaefer-100) preserving the bilateral distribution of channels. The first two gradients of neural dynamics similarity closely resembled those obtained from the unilateral mapping of channels to a higher resolution parcellation. Projection of G1 and G2 to the cortical surface emphasized strong differentiation of neural dynamics between anterior-posterior (G1) and unimodal-transmodal (G2) regions. Source data are provided as a Source Data file.

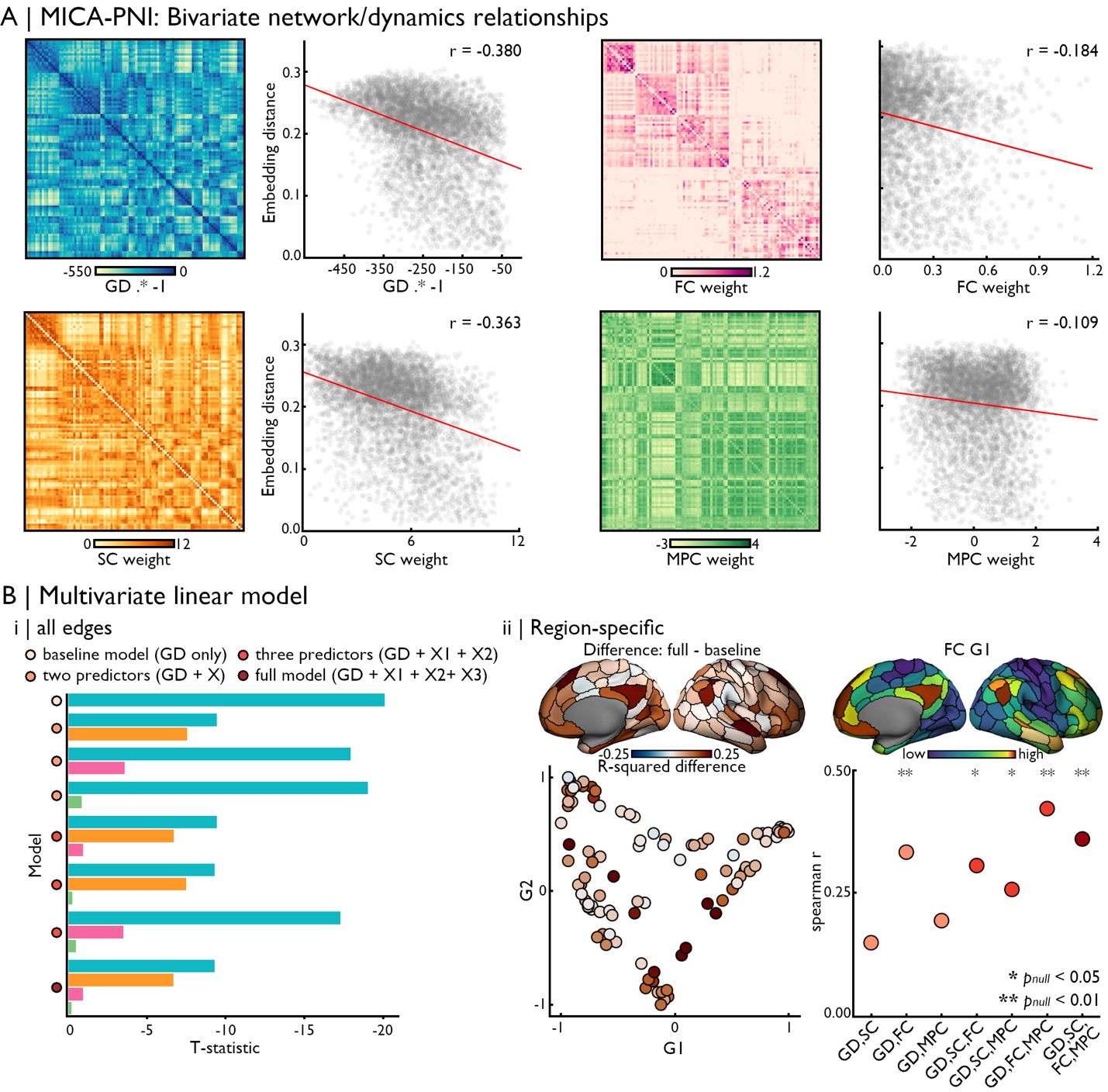

**Figure S5. Multiscale connectomics of neural dynamics at ultra-high fields. (A)** Modality-specific correlations are shown for geodesic distance, structural connectivity, functional connectivity, and microstructural profile similarity. Relative differences across bivariate relationships for each modality followed patterns of findings observed with 3T data. **(B)** Multivariate models with 7T MRI also followed results obtained at 3T, with the strongest contribution of GD when implementing models across all matrix edges. Models implemented at a node level revealed a selective gain in adjusted r^2^ for more complex models including measures of FC and MPC in transmodal cortices. Source data are provided as a Source Data file.

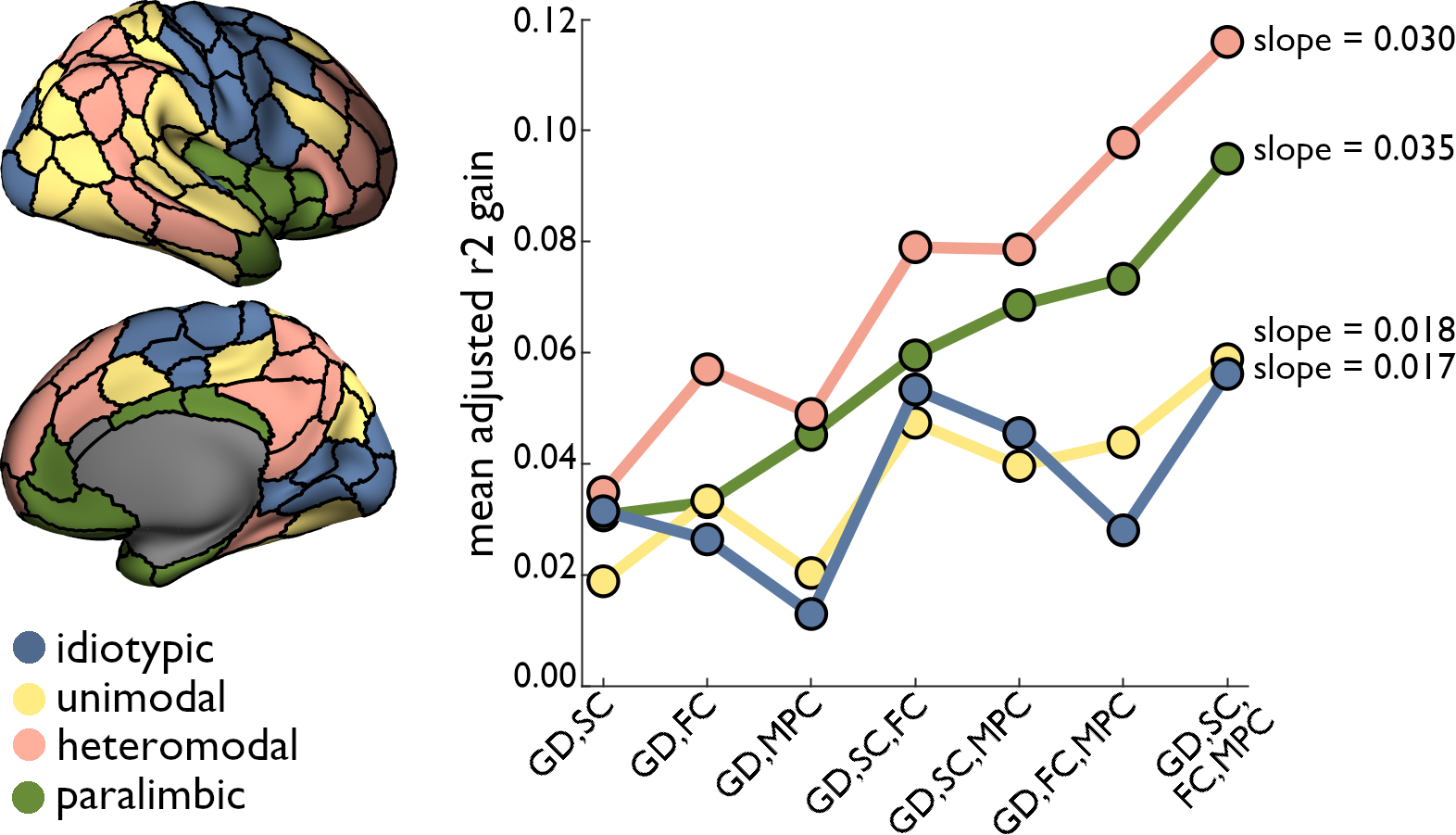

**Figure S6. Multivariate model accuracy gains track the cortical hierarchy.** Parcels were assigned to a functional zone as defined by Mesulam (1998, 2000) based on their overlap in a winner-takes-all fashion (left). Gains in adjusted r^2^ for each multivariate models over the univariate GD model were averaged within these zones (right). Heteromodal and paralimbic cortices selectively benefitted from the inclusion of multimodal predictors over idiotypic and unimodal areas. Slopes were computed via a simple linear regression fit over class-specific mean adjusted r^2^ gain and the number of predictors included in each model. Source data are provided as a Source Data file.

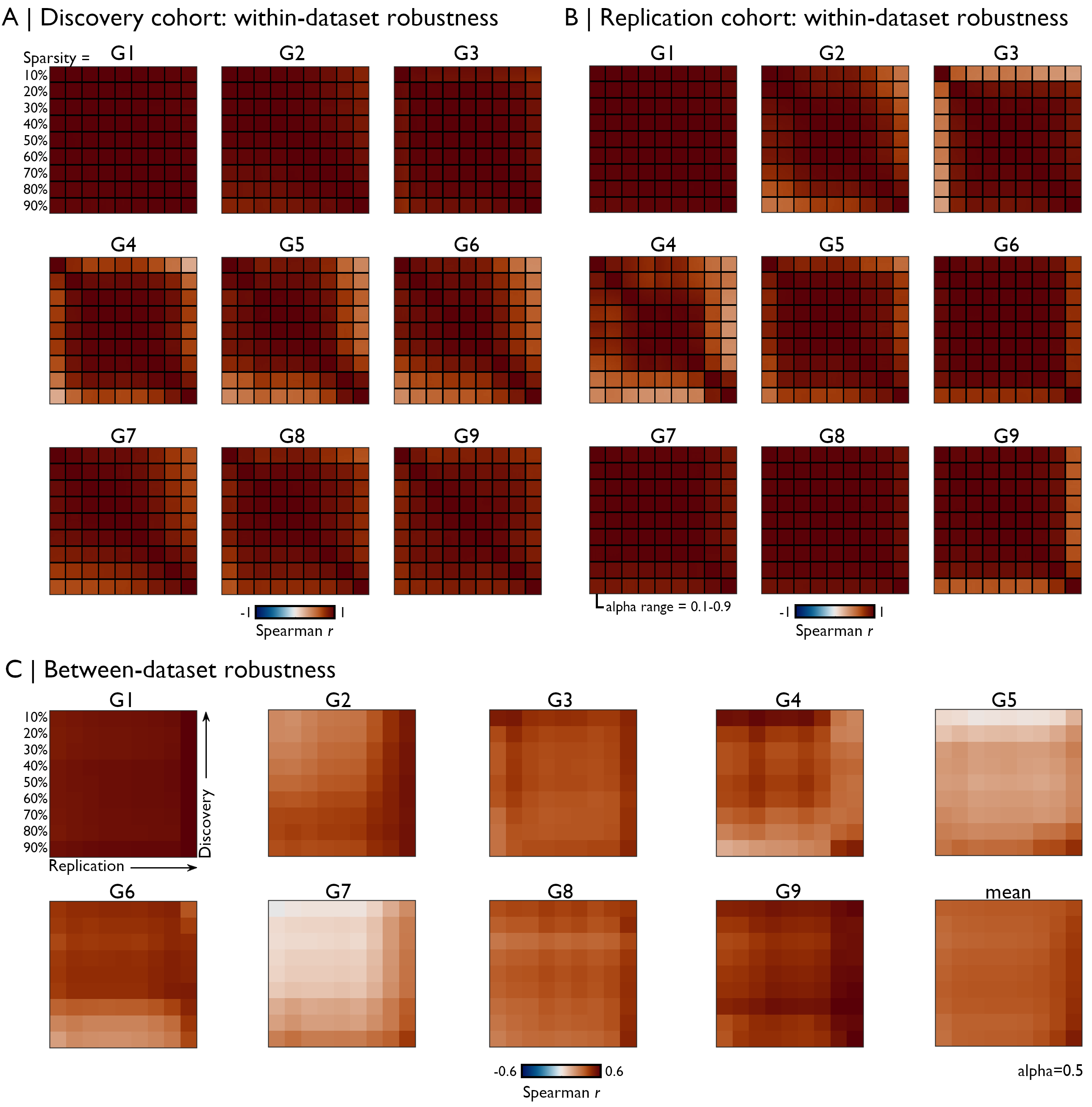

**Figure S7. Robustness to hyperparameter variations within and across datasets.** We systematically cross-correlated each gradient generated from a range of sparsity and alpha parameters within the discovery **(A)** and replication **(B)** cohorts. All gradients were aligned to a common template (sparsity=90%, alpha=0.5, from the discovery cohort) using Procrustes alignment to ensure the consistency of their signs. Small squares within each matrix bound a given sparsity value, and entries within each of these submatrices refer to distinct alpha parameter values. Note that for a given sparsity value, gradients generated from different alpha values were highly similar, hence the relatively homogeneous colouring of each submatrix. **(C)** For a given alpha of 0.5, we assessed the cross-dataset consistency of each gradient over a range of sparsity values. For gradients explaining more variance in the input data, imposing higher matrix sparsity generally resulted in higher reproducibility.

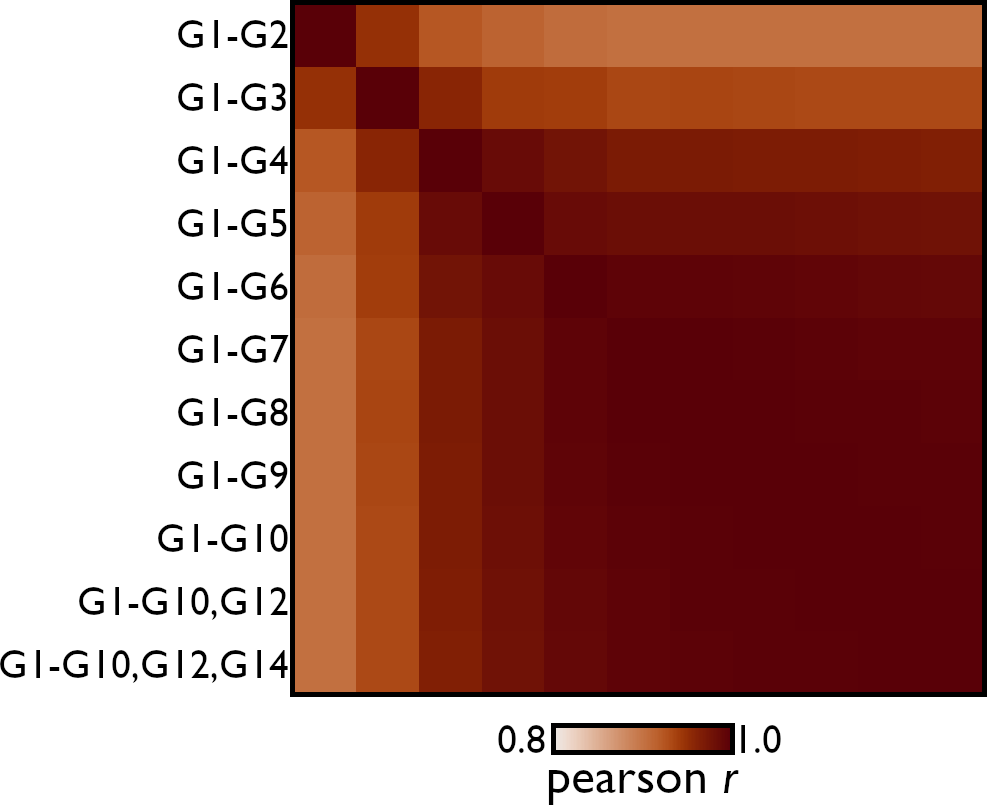

**Figure S8. Robustness to variations in number of gradients included in embedding distance matrix.** We systematically varied the number of gradients included when computing inter-node embedding distances and cross-correlated each matrix (excluding the diagonal). The resulting correlation matrix (min *r* = 0.902; mean *r* = 0.965) shows high reproducibility of the embedding distance matrix when successively including more gradients to compute embedding distances.

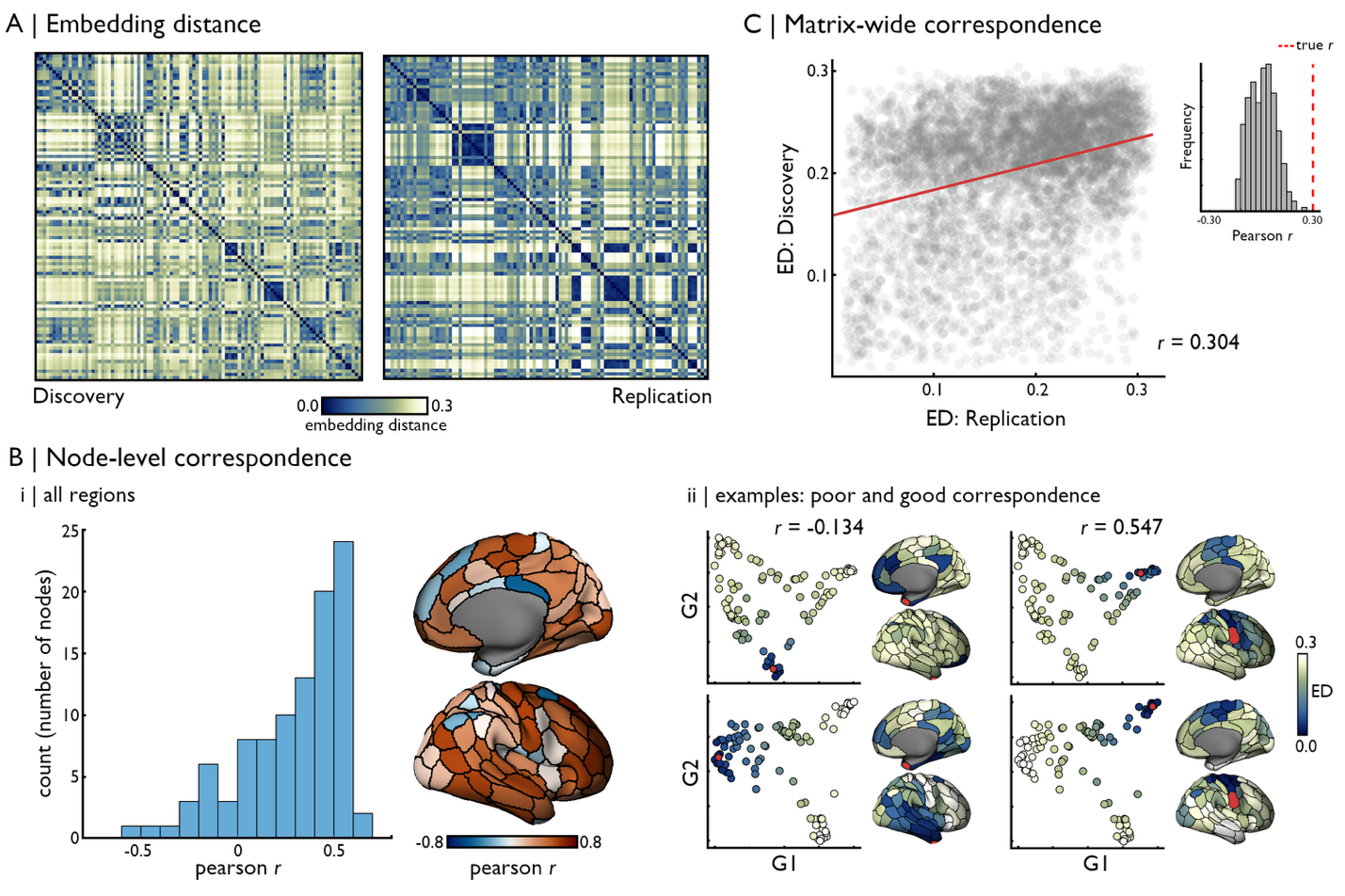

**Figure S9. Replication of embedding distances across Discovery and Replication cohorts. (A)** Embedding distance matrices for each cohort. **(B)** Cross-correlating corresponding rows of each matrix showed variable correspondence of inter-node distances, with the distribution of correlation coefficients (Pearson’s *r*) shown in a histogram. Correlation coefficients for each region are also displayed on the cortical surface, alongside example seeds with poor correspondence (10^th^ percentile; temporal pole) and good correspondence (90^th^ percentile; central sulcus) across cohorts. Selected seeds are shown in red. **(C)** Correlating the upper triangular part of each embedding distance matrix showed a moderate (*r* = 0.304; *p*<0.001) correlation across cohorts. Statistical significance was determined by comparing empirically observed correlations to those obtained with surrogate matrices in which each row had been shuffled using spatial autocorrelation-preserving null models (as in **Figure 2** of the main manuscript). Source data are provided as a Source Data file.

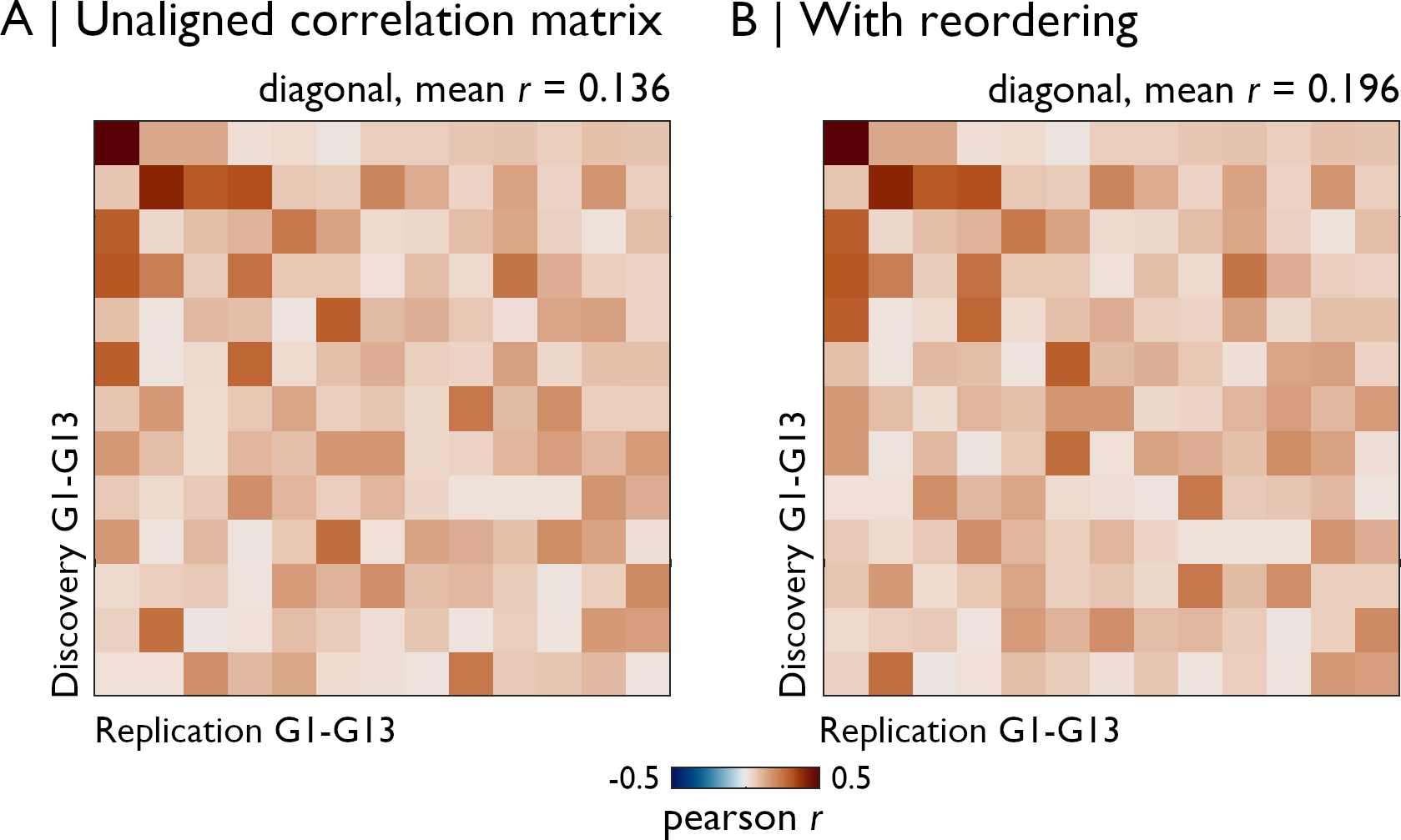

**Figure S10. Gradient alignment across Discovery and Replication cohorts. (A)** The absolute value of Spearman *r* of gradients across cohort (without alignment). We show the absolute value of correlation coefficients as signs of eigenvectors can be flipped across datasets. **(B)** Reordering gradients in the Replication cohort to minimize their distance with gradients of the Discovery cohort could improve Spearman *r* values across the correlation matrix diagonal.

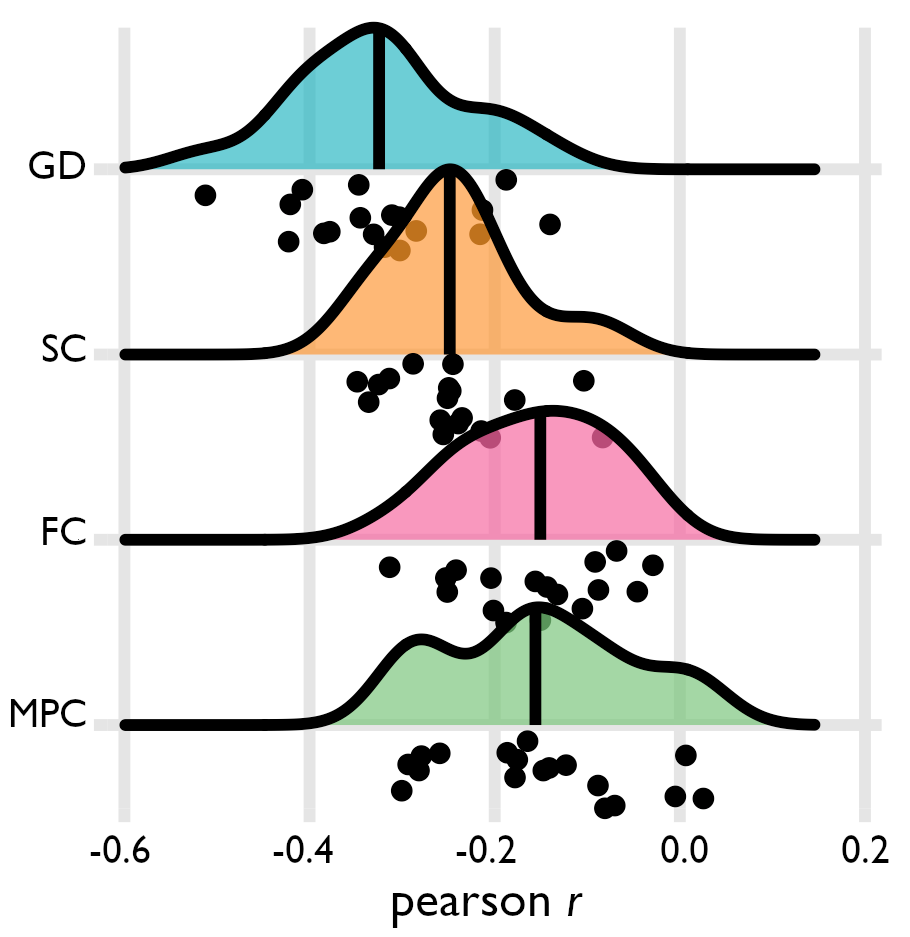

**Figure S11. Individual-level correspondence of iEEG-MRI associations in Replication cohort.** We computed patient-specific gradients from covered regions and computed the embedding distance between available nodes for gradients explaining up to 50% in the input data. Bilateral implantations were mapped to the right hemisphere (n=4), left-sided implantations were flipped to the right hemisphere (n=5), and right hemisphere implantation were analysed without flipping (n=9). For each patient, embedding distances were correlated with MRI features of corresponding nodes, and resulting correlation coefficients (Pearson *r*) are represented as black markers. Each black dot represents a single patient, with the group distribution shown in ridge plots for each modality (black line: median). Replicating group-level findings in the Discovery cohort, similarity of neural dynamics in the cortex were most strongly correlated with geodesic distance (GD; median *r*=-0.325), followed by structural connectivity (SC; median *r* = -0.249), and lower associations with functional connectivity (median *r* = -0.151) and microstructural profile covariance (median *r* = -0.156). Source data are provided as a Source Data file.

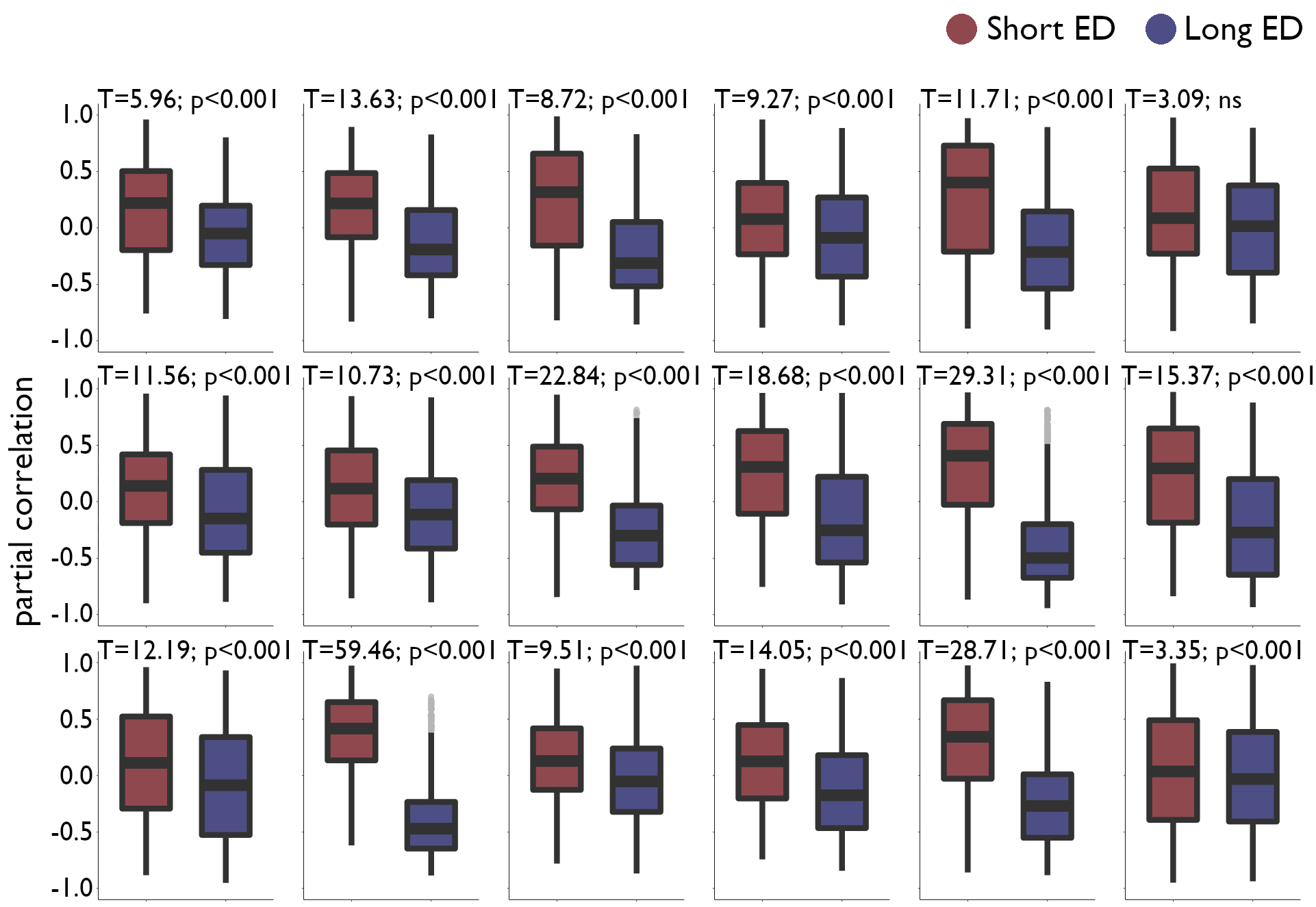

**Figure S12. Similarity of PSDs across short and long embedding distances**. The group-level gradient space was regenerated while iteratively excluding each subject, and we calculated embedding distances in each group-level space. For the left-out subject at each iteration, we evaluated if the proximity of sampled nodes in the embedding space was related to the similarity (partial correlation of PSDs, while controlling for the within-subject average) of corresponding PSDs. We specifically compared partial correlation values at short embedding distances (embedding distance <20^th^ percentile) versus long embedding distances (embedding distance >80^th^ percentile) for each patient. In all patients (each represented by their own boxplot), PSDs were more highly correlated across nodes situated closer in the embedding space computed from the remaining 17 patients, although the strength of this effect varied across individuals (mean T-statistic=16.004; S.D.=13.152). All p-values are FDR-corrected. Source data are provided as a Source Data file.
